# The AMA1-RON complex drives *Plasmodium* sporozoite invasion in the mosquito and mammalian hosts

**DOI:** 10.1101/2022.01.04.474787

**Authors:** Priyanka Fernandes, Manon Loubens, Rémi Le Borgne, Carine Marinach, Beatrice Ardin, Sylvie Briquet, Laetitia Vincensini, Soumia Hamada, Bénédicte Hoareau-Coudert, Jean-Marc Verbavatz, Allon Weiner, Olivier Silvie

**Author notes:** These authors contributed equally to the work (Priyanka Fernandes, Manon Loubens).

## Abstract

*Plasmodium* sporozoites that are transmitted by blood-feeding female *Anopheles* mosquitoes invade hepatocytes for an initial round of intracellular replication, leading to the release of merozoites that invade and multiply within red blood cells. Sporozoites and merozoites share a number of proteins that are expressed by both stages, including the Apical Membrane Antigen 1 (AMA1) and the Rhoptry Neck Proteins (RONs). Although AMA1 and RONs are essential for merozoite invasion of erythrocytes during asexual blood stage replication of the parasite, their function in sporozoites is still unclear. Here we show that AMA1 interacts with RONs in mature sporozoites. By using DiCre-mediated conditional gene deletion in *P. berghei*, we demonstrate that loss of AMA1, RON2 or RON4 in sporozoites impairs colonization of the mosquito salivary glands and invasion of mammalian hepatocytes, without affecting transcellular parasite migration. Our data establish that AMA1 and RONs facilitate host cell invasion across *Plasmodium* invasive stages, and suggest that sporozoites use the AMA1-RON complex to safely enter the mosquito salivary glands without causing cell damage, to ensure successful parasite transmission. These results open up the possibility of targeting the AMA1-RON complex for transmission-blocking antimalarial strategies.

## Introduction

Host cell invasion is an obligatory step in the *Plasmodium* life cycle. There are several invasive stages of *Plasmodium*, each equipped with its own set of specialised secretory organelles and proteins that facilitate invasion into or through host cells. Invasive stages of Apicomplexa typically invade target host cells actively by gliding through a structure known as the moving junction (MJ), which consists of a circumferential zone of close apposition of parasite and host cell membranes. Studies with *Toxoplasma gondii* tachyzoites and *Plasmodium falciparum* merozoites have shown that formation of the MJ involves the export of rhoptry neck proteins RONs into the host cell, where RON2 is inserted into the host cell membrane and serves as a receptor for the Apical Membrane Antigen 1 (AMA1), that is secreted from the micronemes onto the surface of the parasite (Besteiro et al., 2011; Cowman et al., 2017; Frénal et al., 2017). Formation of the MJ is associated with active penetration inside the parasitophorous vacuole (PV), which is essential for further development and replication of the parasite.

Although the AMA1-RON2 interaction seems to be conserved across the phylum of Apicomplexa, its role in *Plasmodium* sporozoites is controversial. *Plasmodium* sporozoites express AMA1 and the RON proteins RON2, RON4 and RON5 (Giovannini et al., 2011; Hamada et al., 2021; Lindner et al., 2013; Silvie et al., 2004; Swearingen et al., 2017; Tokunaga et al., 2019; Tufet-Bayona et al., 2009). Two studies reported that AMA1 is not essential for development in the mosquito and during hepatocyte invasion in *P. berghei*, while RON4 in contrast was shown to be essential for hepatocyte invasion, suggesting independent roles for AMA1 and RON proteins in sporozoites (Bargieri et al., 2013; Giovannini et al., 2011). However, both polyclonal antibodies against AMA1 (Silvie et al., 2004) and the R1 peptide inhibitor of AMA1 (Harris et al., 2005), effectively reduced hepatocyte invasion by *P. falciparum* sporozoites (Yang et al., 2017). More recently, a promoter swap strategy was employed to knockdown RONs in *P. berghei* sporozoites, uncovering an unexpected role of these proteins during invasion of the mosquito salivary glands (Ishino et al., 2019; Nozaki et al., 2020). Owing to these conflicting data, the precise role of AMA1 and RONs in *Plasmodium* sporozoites is uncertain.

As conventional reverse genetics cannot be used to target AMA1 and RONs, due to their essential nature in asexual blood stages, previous studies relied on conditional approaches such as the Flippase (FLP)/Flp recombination target (FRT) system or promoter swap strategies to target these genes. The rapamycin inducible DiCre recombinase system has recently emerged as a potent method of gene inactivation in different developmental stages of the parasite life cycle, in *P. falciparum* (Tibúrcio et al., 2019) and *P. berghei* (Fernandes et al., 2020). We recently described a fluorescent DiCre-expressing parasite line in *P. berghei* and showed that efficient and complete gene excision can be induced in asexual blood stages and also sporozoites (Fernandes et al., 2020). In this study, we used the DiCre system to achieve conditional deletion of *ama1*, *ron2* and *ron4* genes in *P. berghei* sporozoites. Our data reveal that sporozoites rely on AMA1 and RONs to invade salivary glands in the mosquito and hepatocytes in the mammalian host, implying a conserved feature of the invasion process across invasive stages of *Plasmodium*.

## Results

### Deletion of *ama1* 3’UTR is not sufficient to abrogate AMA1 expression in *P. berghei*

To ablate AMA1 protein expression in *P. berghei*, we first decided to conditionally delete the 3’ untranslated region (UTR) of *ama1* using the DiCre method, as previously reported with the FLP/FRT system (Giovannini et al., 2011). We floxed the 3’UTR of *ama1*, together with a GFP and an hDHFR marker, to generate the *ama1*Δutr parasite line in the mCherry-expressing PbDiCre parasite background (Fernandes et al., 2020) (**Figure 1A** and **Figure 1-Supplement 1**). To exclude any unspecific effects arising from modification of the *ama1* locus, we also generated a control parasite line (*ama1*Con) where we introduced the LoxN sites downstream of the 3’ UTR (**Figure 1B** and **Figure 1-Supplement 2**). After transfection and selection with pyrimethamine, pure populations of recombinant parasites were sorted by flow cytometry and genotyped by PCR to confirm correct genomic integration of the constructs and to exclude the presence of any residual unmodified PbDiCre parasites (**Figure 1-Supplements 1** and **2**).

**Figure 1.**
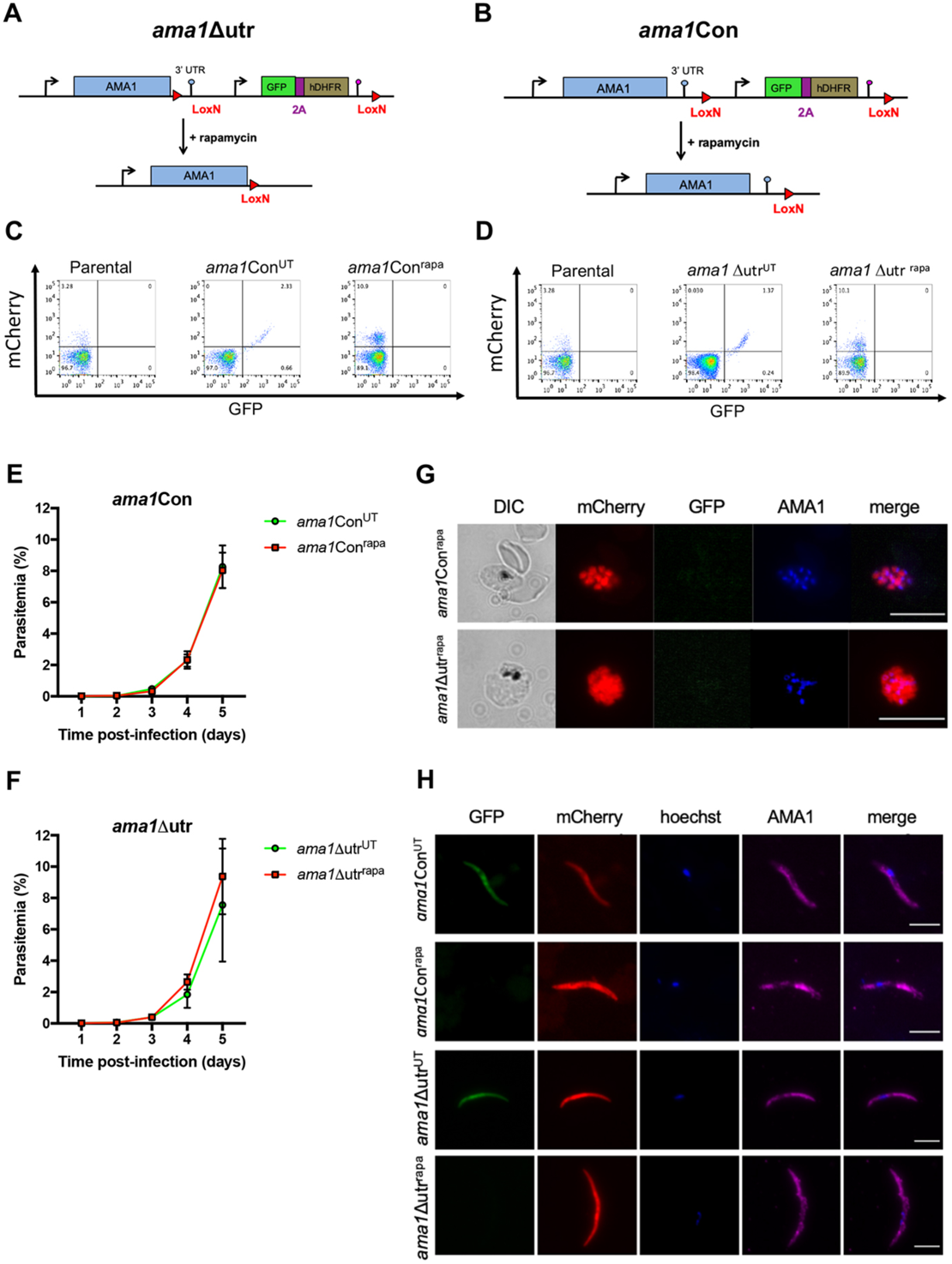
Deletion of the 3’ UTR of *ama1* has no phenotypical impact in *P. berghei*. **A-B.** Strategy to generate *ama1*Δutr (A) and *ama1*Con (B) parasites by modification of the wild type *ama1* locus in PbDiCre parasites. **C-D.** Flow cytometry analysis of PbDiCre (parental), *ama1*Con (C) and *ama1*Δutr (D) untreated (^UT^) or rapamycin-treated (^rapa^) blood stage parasites. **E-F.** Blood stage growth of treated (^rapa^) and untreated (^UT^) *ama1*Con (E) or *ama*1Δutr (F) parasites. Rapamycin was administered at day 2. The graphs represent the parasitaemia (mean +/- SEM) in groups of 3 mice. **G.** Immunofluorescence staining of rapamycin-treated *ama1*Con and *ama1*Δutr blood stage schizonts with anti-AMA1 antibodies (blue). Scale bar = 10 µm. **H.** Immunofluorescence images of rapamycin-treated *ama1*Con and *ama1*Δutr sporozoites after staining with anti-AMA1 antibodies (magenta). Scale bar = 5 µm.

We next analyzed the effects of rapamycin on *ama1*Con and *ama1*Δutr parasites during blood stage growth, by quantifying the percentage of excised (mCherry^+^/GFP^−^) and non-excised (mCherry^+^/GFP^+^) parasites by flow cytometry (**Figure 1C** and **1D**). In the *ama1*Con infected group, rapamycin treatment induced complete excision of the floxed GFP cassette (*ama1*Con^rapa^) (**Figure 1C**), which, as expected, had no significant effect on parasite growth and multiplication in the blood, which was comparable to the untreated group (*ama1*Con^UT^) (**Figure 1E**). Excision of the GFP cassette was also confirmed by genotyping PCR (**Figure 1-Supplement 2**). Surprisingly, rapamycin treatment of the *ama1*Δutr infected group (*ama1*Δutr^rapa^) also had no effect on both parasite growth and multiplication in the blood (**Figure 1F**), despite efficient DNA excision based on disappearance of the GFP cassette after rapamycin treatment (**Figure 1D**). Genotyping of mCherry^+^/GFP^−^ parasites by PCR and sequencing of the locus after excision confirmed that the 3’UTR had been excised in *ama1*Δutr^rapa^ parasites, excluding any contamination with parental PbDiCre (**Figure 1-Supplement 2**).

To explore whether *ama1*Δutr^rapa^ parasites had adapted despite losing AMA1 expression, we fixed *ama1*Con^rapa^ and *ama1*Δutr^rapa^ blood-stage schizonts and incubated them with anti-AMA1 antibodies for immunofluorescence staining. Intriguingly, we observed AMA1 expression in both *ama1*Con^rapa^ and *ama1*Δutr^rapa^ merozoites (**Figure 1G**) implying that deletion of the *ama1* 3’UTR alone is not sufficient to abrogate expression of the protein in merozoites. We further analysed the impact of 3’UTR deletion on AMA1 expression in sporozoites. For this purpose, *ama1*Con and *ama1*Δutr parasites were treated with rapamycin or left untreated and then transmitted to mosquitoes, as described previously (Fernandes et al., 2020). Deletion of the *ama1* 3’UTR in *ama1*Δutr^rapa^ parasites had no impact on oocyst formation in the midgut or sporozoite invasion of salivary glands, which were comparable to *ama1*Δutr^UT^ and both *ama1*Con^UT^ and *ama1*Con^rapa^ parasites (**Figure 1-Supplement 3**). Consistent with AMA1 protein expression in *ama1*Δutr^rapa^ merozoites, we also detected AMA1 protein in *ama1*Δutr^rapa^ salivary gland sporozoites by immunofluorescence, similar to *ama1*Con^rapa^ parasites (**Figure 1H**). We conclude from these data that deletion of the 3’UTR of *ama1* is not sufficient to abrogate AMA1 protein expression and cause phenotypical changes in *P. berghei* merozoites and sporozoites.

### Complete conditional gene deletion of *ama1* in *P. berghei*

Since deletion of the 3’UTR was insufficient to deplete AMA1, we decided to delete the full-length *ama1* gene, by placing LoxN sites both upstream and downstream of the gene (**Figure 2A**). One intrinsic feature of the Cre Lox system is the retention of a Lox site following recombination. We therefore reused *ama1*Con^rapa^ parasites, which contained a single LoxN site downstream of *ama1* 3’UTR, and transfected these parasites with the *ama1*cKO construct designed to introduce a second LoxN site upstream of the *ama1* gene, together with a GFP-hDHFR cassette (**Figure 2A** and **Figure 2-Supplement 1**). Following transfection, the resulting *ama1*cKO parasites were sorted by FACS and genotyped to confirm correct integration of the construct into the genome and verify the absence of any residual unmodified *ama1*Con parasites (**Figure 2-Supplement 1**). We then evaluated the effect of rapamycin treatment on blood-stage growth of *ama1*cKO parasites, by injecting mice with 10^6^ pRBCs and treating them with a single oral dose of rapamycin. In contrast to untreated *ama1*cKO (*ama1*cKO^UT^) infected mice, parasite growth was abrogated in mice upon rapamycin exposure (*ama1*cKO^rapa^) (**Figure 2B**), thus confirming efficient gene deletion and the essential role of AMA1 in merozoite invasion and parasite survival in the blood.

**Figure 2.**
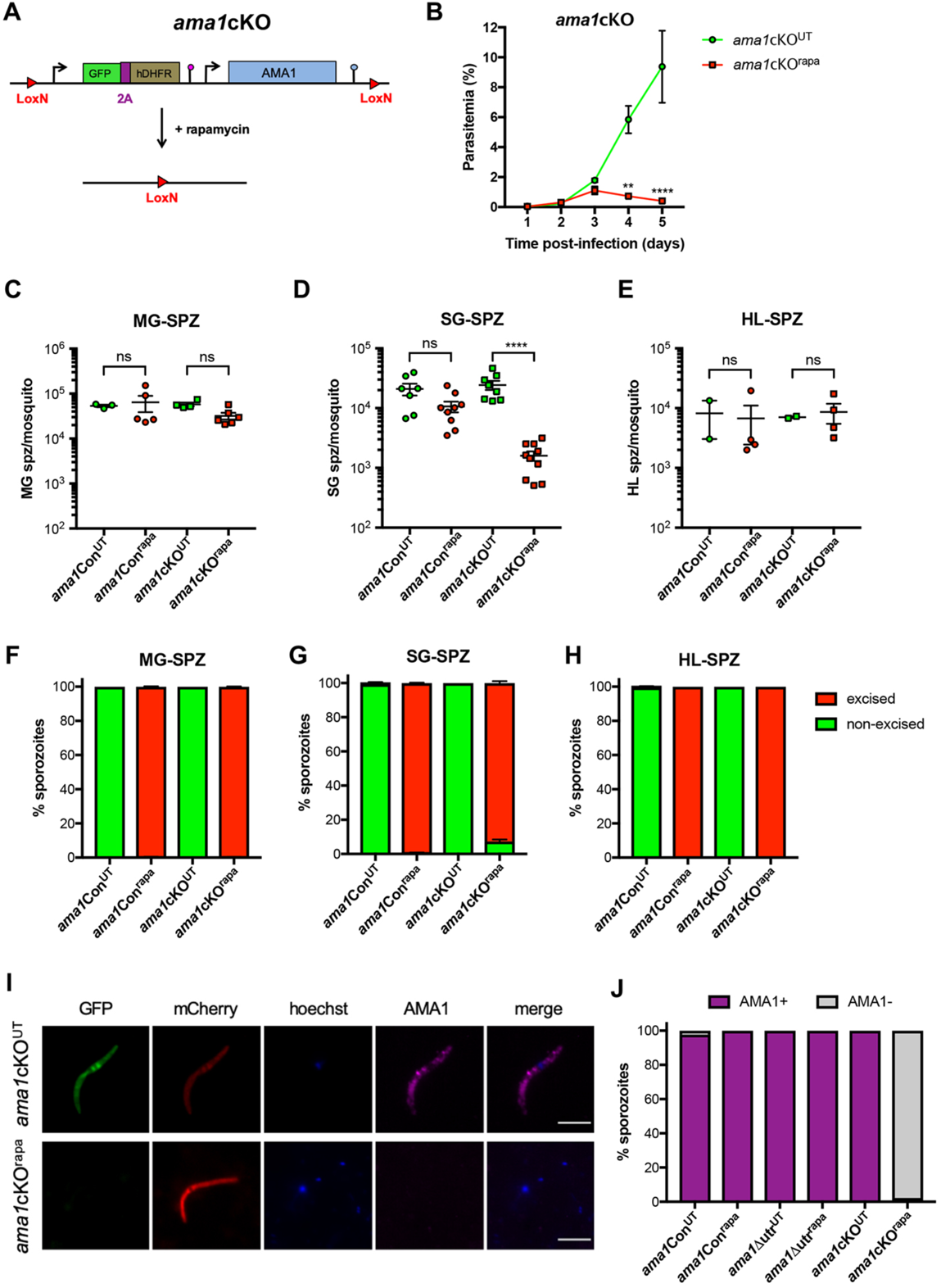
AMA1 is required during *P. berghei* invasion of mosquito salivary glands. **A.** Strategy to generate *ama1*cKO parasites by modification of the *ama1* locus in rapamycin-treated *ama1*Con parasites. **B.** Blood stage growth of treated (^rapa^) and untreated (^UT^) *ama1*cKO parasites. The graph represents the parasitaemia (mean +/- SEM) in groups of 3 mice. Rapamycin was administered at day 2. **, p < 0.01; ****, p < 0.0001 (Two-way ANOVA). **C-E.** Quantification of midgut sporozoites (MG-SPZ, C), salivary gland sporozoites (SG-SPZ, D) or haemolymph sporozoites (HL-SPZ, E) isolated from mosquitoes infected with untreated (^UT^) or rapamycin treated (^rapa^) *ama1*Con and *ama1*cKO parasites. The graphs show the number of sporozoites per female mosquito (mean +/- SEM). Each dot represents the mean value obtained in independent experiments after dissection of 30-50 mosquitoes (MG, HL) or 50-70 mosquitoes (SG), respectively. Ns, non-significant; ****, p < 0.0001 (One-way ANOVA followed by Tukey’s multiple comparisons test). **F-H.** Quantification of excised (mCherry^+^/GFP^−^, red) and non-excised (mCherry^+^/GFP^+^, green) midgut sporozoites (MG-SPZ, F), salivary gland sporozoites (SG-SPZ, G) or haemolymph sporozoites (HL-SPZ, H) isolated from mosquitoes infected with untreated (^UT^) or rapamycin treated (^rapa^) *ama1*Con and *ama1*cKO parasites. **I.** Immunofluorescence imaging of untreated (^UT^) and rapamycin-treated (^rapa^) *ama1*cKO salivary gland sporozoites after staining with anti-AMA1 antibodies (magenta). Scale bar = 5 µm. **J.** Quantification of AMA1-positive and AMA1-negative sporozoites among untreated (^UT^) or rapamycin-treated (^rapa^) *ama1*Con, *ama1*Δutr and *ama1*cKO sporozoites, as assessed by microscopy.

### AMA1 is required for sporozoite invasion of the mosquito salivary glands

In order to determine the function of AMA1 in sporozoites, we transmitted *ama1*cKO^UT^ and *ama1*cKO^rapa^ parasites to mosquitoes, 24 hours after rapamycin treatment. In parallel, mosquitoes were fed with *ama1*Con^UT^ and *ama1*Con^rapa^ parasites as a reference line. Both *ama1*cKO^UT^ and *ama1*cKO^rapa^ parasites were capable of colonising the mosquito midgut (**Figure2-Supplement 2**), comparable to *ama1*Con parasites (**Figure 1-Supplement 3**). Despite no difference in the levels of exflagellation between all four parasite lines, we observed a slight reduction in the number of midgut sporozoites for *ama1*cKO^rapa^ parasites, which however was not statistically significant (**Figure 2C**). Importantly, quantification of the percentage of excised (mCherry^+^/GFP^−^) and non-excised (mCherry^+^/GFP^+^) parasites revealed close to 100% gene excision in sporozoites isolated from the midguts of mosquitoes infected with *ama1*Con^rapa^ and *ama1*cKO^rapa^ parasites (**Figure 2F**).

In the next step, we quantified sporozoites isolated from the salivary glands of infected mosquitoes and observed no difference between mosquitoes infected with *ama1*Con^UT^ or *ama1*cKO^UT^ parasites (**Figure 2D**). In sharp contrast, the number of salivary gland sporozoites isolated from *ama1*cKO^rapa^ infected mosquitoes was severely reduced as compared to untreated parasites (**Figure 2D** and **Figure 2-Supplement 2**). As expected, we could only observe mCherry^+^/GFP^+^ (non-excised) salivary gland sporozoites in *ama1*Con^UT^ and *ama1*cKO^UT^ parasites, while *ama1*Con^rapa^ and *ama1*cKO^rapa^ sporozoites were mCherry^+^/GFP^−^ (excised) (**Figure 2G**). Interestingly, a small proportion (<10%) of *ama1*cKO^rapa^ salivary gland sporozoites were mCherry^+^/GFP^+^ (non-excised), suggesting an enrichment of sporozoites harbouring an intact *ama1* gene, in the salivary glands of infected mosquitoes (**Figure 2G**).

In order to determine if a defect in egress from oocysts or invasion of the salivary glands was the reason behind the reduction in *ama1*cKO^rapa^ salivary gland sporozoite numbers, we quantified haemolymph sporozoites from infected mosquitoes at day 14 post infection. There was no significant difference between the numbers of haemolymph sporozoites isolated from *ama1*Con and *ama1*cKO infected mosquitoes before or after rapamycin treatment (**Figure 2E**). Using microscopy, we could only see non-excised (mCherry^+^/GFP^+^) haemolymph sporozoites for *ama1*Con^UT^ and *ama1*cKO^UT^ infected mosquitoes, while all *ama1*Con^rapa^ and *ama1*cKO^rapa^ haemolymph sporozoites were excised (mCherry^+^/GFP^−^) (**Figure 2H**). The absence of a defect in egress from oocysts was also documented by microscopy imaging of the abdomen of infected mosquitoes, where scavenging of circulating sporozoites following egress results in bright red fluorescence of pericardial cellular structures (**Figure 2-Supplement 3**). A similar percentage of mosquitoes displayed mCherry-labelled pericardial cells between untreated and rapamycin treated *ama1*Con and *ama1*cKO infected mosquitoes, confirming that loss of AMA1 expression in sporozoites does not affect sporozoite egress from oocysts (**Figure 2-Supplement 3**).

Lastly, we verified the loss of AMA1 expression in sporozoites by immunofluorescence imaging of salivary gland sporozoites using anti-AMA1 antibodies. The micronemal pattern of AMA1 expression was consistent between *ama1*Con^UT^ and *ama1*Con^rapa^ sporozoites, as well as *ama1*cKO^UT^ sporozoites (**Figure 1H and 2I**). However, no AMA1 was detected in *ama1*cKO^rapa^ sporozoites, indicating the loss of AMA1 (**Figure 2I**). Quantification of AMA1 expression showed that all sporozoites from *ama1*Con^UT^, *ama1*Con^rapa^ and *ama1*cKO^UT^, as well as *ama1*Δutr^UT^ and *ama1*Δutr^rapa^ parasites, expressed AMA1 (**Figure 2J**). In contrast, >90% of the sporozoites isolated from *ama1*cKO^rapa^ infected mosquitoes lacked AMA1, confirming successful gene deletion and protein depletion (**Figure 2J**). Overall, our results demonstrate that loss of AMA1 expression in sporozoites impairs invasion of the mosquito salivary glands, without affecting development or egress from oocysts.

### AMA1 is required for efficient sporozoite invasion of hepatocytes

In the next step, we tested if *ama1*cKO^rapa^ sporozoites could infect hepatocytes. AMA1 was previously suggested to be implicated in cell traversal of *P. falciparum* sporozoites (Yang et al., 2017). Hence we first verified if *ama1*cKO^rapa^ sporozoites also displayed a similar defect in cell traversal *in vitro,* using a dextran assay as previously described (Mota et al., 2001). Quantification of dextran-positive cells indicated that cell traversal was comparable between *ama1*con^rapa^ and *ama1*cKO^rapa^ sporozoites, implying that both sporozoite motility and cell traversal were unaffected by excision of *ama1* (**Figure 3A**). We then infected HepG2 cell cultures with sporozoites isolated from mosquitoes previously fed with rapamycin-treated or untreated *ama1*Con and *ama1*cKO parasites. We quantified infected cells, containing exo-erythrocytic forms (EEFs), at 24 h post infection by flow cytometry and fluorescence microscopy. We observed a minor but non-significant reduction in the number of EEFs for *ama1*Con^rapa^ parasites (**Figure 3B**) compared to *ama1*Con^UT^. In contrast, the number of EEFs obtained from hepatocytes infected with *ama1*cKO^rapa^ sporozoites was significantly reduced (**Figure 3B**). As expected, non-excised (mCherry^+^/GFP^+^) parasites comprised the majority of EEFs quantified for *ama1*Con^UT^ and *ama1*cKO^UT^ parasites (**Figure 3C**). Conversely, excised (mCherry^+^/GFP^−^) EEFs were predominantly observed for both *ama1*Con^rapa^ and *ama1*cKO^rapa^ infected hepatocytes. However, an enrichment of non-excised (mCherry^+^/GFP^+^) EEFs was observed in *ama1*cKO^rapa^ infected hepatocytes (**Figure 3C**), similar to what was observed for the salivary glands of mosquitoes infected with *ama1*Con^rapa^ parasites (**Figure 2G**). Importantly, we could not observe any obvious defect in developmental size or morphology for all four parasite lines, by fluorescence microscopy (**Figure 3D**). Finally, UIS4 staining of the PV membrane confirmed that mCherry^+^/GFP^−^ (excised) *ama1*cKO^rapa^ sporozoites could form a PV *in vitro*, similar to *ama1*Con^rapa^ EEFs (**Figure 3E**), implying that in the absence of AMA1, sporozoites conserve a residual capacity to productively invade host cells.

**Figure 3.**
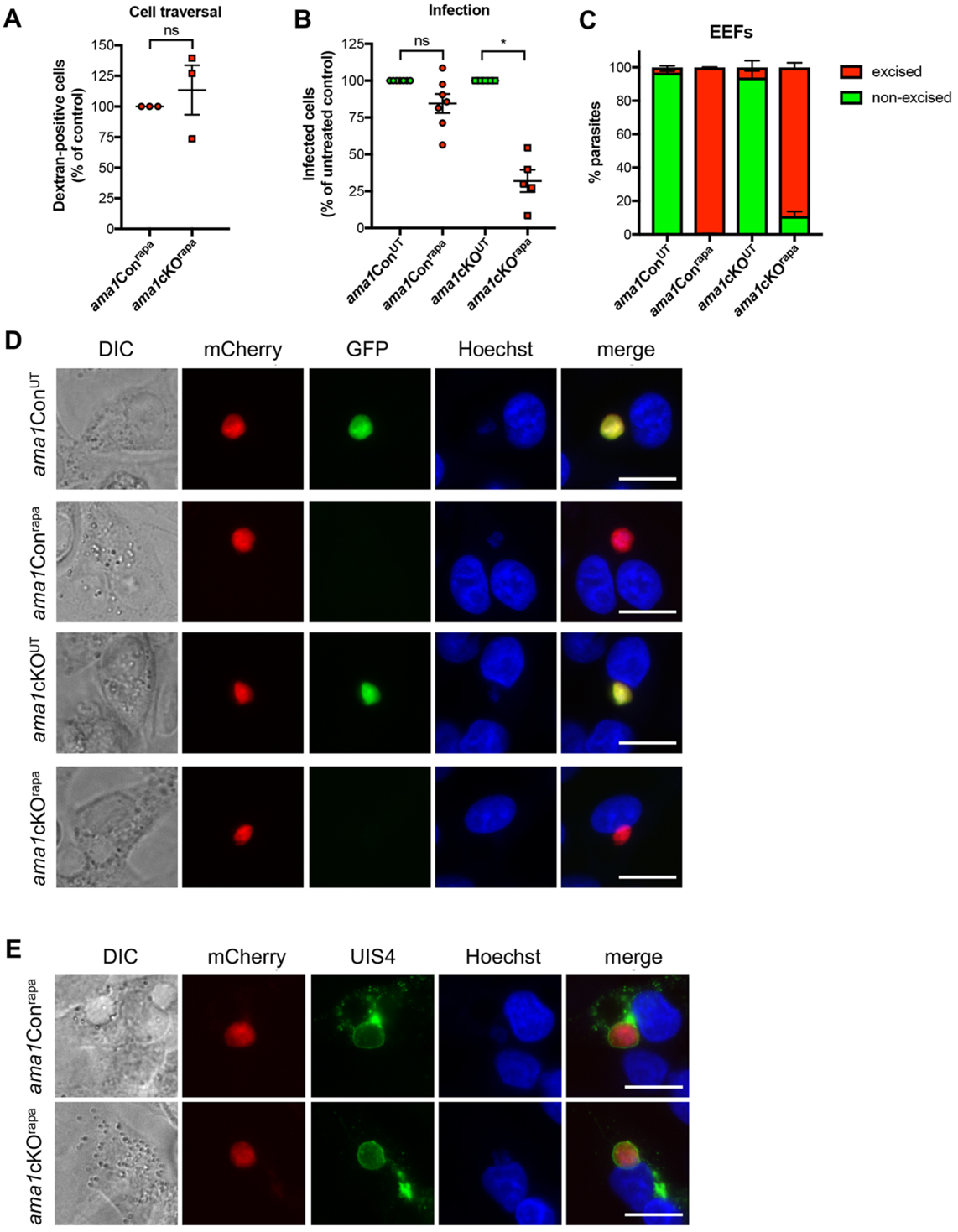
Sporozoite AMA1 is required for efficient infection of mammalian cells. **A.** Quantification of sporozoite cell traversal activity (% of dextran-positive cells) in rapamycin-treated *ama1*Con and *ama1cKO* parasites. The values for rapamycin-treated *ama1cKO* parasites are represented as percentage of the rapamycin-treated *ama1*Con parasites (mean +/- SEM of three independent experiments). Each data point is the mean of five technical replicates. Ns, non-significant (Two-tailed ratio paired t test). **B.** Quantification of EEFs development *in vitro,* done by flow cytometry or microscopy analysis of HepG2 cells infected with sporozoites isolated from either untreated (^UT^) or rapamycin-treated (^rapa^) *ama1*Con and *ama1cKO* infected mosquitoes. The data for rapamycin-treated *ama1*Con and *ama1cKO* parasites are represented as percentage of the respective untreated parasites (mean +/- SEM). Each data point is the mean of three technical replicates in one experiment. Ns, non-significant; *, p < 0.05 (Two-tailed ratio paired t test). **C.** Quantification of excised (mCherry^+^/GFP^−^, red) and non-excised (mCherry^+^/GFP^+^, green) EEF populations for untreated and treated *ama1*Con and *ama1cKO* parasites. **D.** Fluorescence microscopy of EEF development (24h p.i.) *in vitro,* in HepG2 cells infected with salivary gland sporozoites from untreated (^UT^) or rapamycin-treated (^rapa^) *ama1*Con and *ama1cKO* parasites. Scale bar = 10 µm. **E.** Immunofluorescence imaging of mCherry^+^/GFP^−^ (excised) rapamycin-treated (^rapa^) *ama1*Con and *ama1*cKO EEFs after staining with anti-UIS4 antibodies (green). Scale bar = 10 µm.

### RON2 and RON4 interact with AMA1 in sporozoites and are required for host cell invasion

Merozoite AMA1 interacts with RON proteins for invasion of erythrocytes (Lamarque et al., 2011; Richard et al., 2010; Srinivasan et al., 2011). In order to investigate whether similar protein interactions also occur in sporozoites, we performed RFP-trap immunoprecipitation experiments using lysates from transgenic sporozoites expressing RON4 fused to mCherry. RON4, RON2, RON5 and AMA1 were the main proteins identified by mass spectrometry among co-precipitated proteins, showing that AMA1-RON interactions are conserved in salivary gland sporozoites (**Table S1**). We decided to focus on RON2 and RON4 and generated conditional mutants, using a two-step strategy to introduce LoxN sites upstream and downstream of the genes in PbDiCre parasites (**Figure 4A** and **Figure 4-Supplements 1** and **2**). Clonal populations of *ron2*cKO and *ron4*cKO parasites were obtained after pyrimethamine selection and FACS sorting, and verified by genotyping PCR (**Figure 4-Supplements 1** and **2**). In agreement with an essential role for RON2 and RON4 in the blood, rapamycin-induced gene excision reduced blood-stage growth in *ron2*cKO^rapa^ and *ron4*cKO^rapa^ infected mice (**Figure 4B** and **4C**).

**Figure 4.**
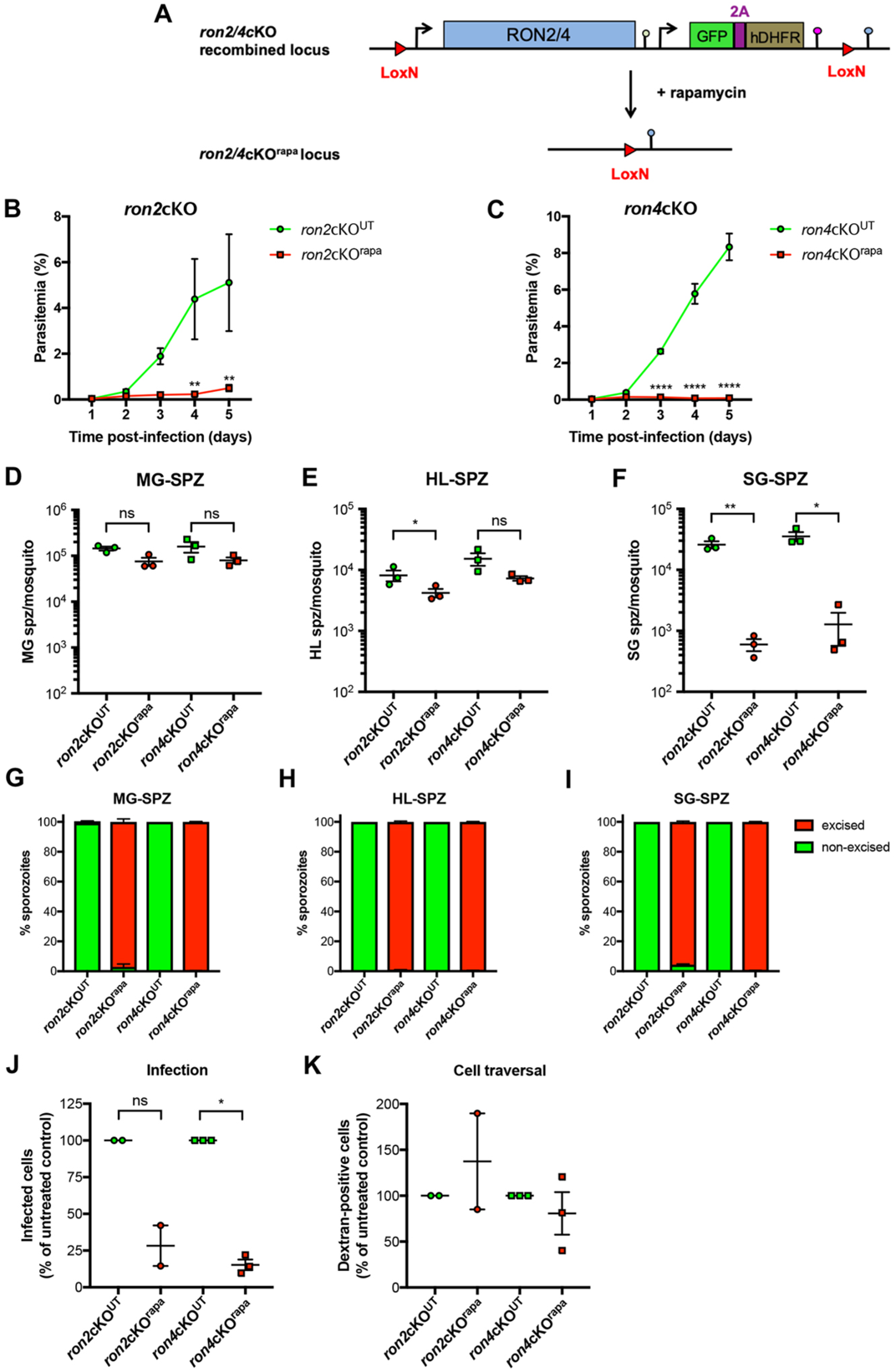
RON2 and RON4 are required for sporozoite invasion in the mosquito and mammalian hosts. **A.** Strategy to generate *ron2*cKO and *ron4*cKO parasites in the PbDiCre line. **B-C.** Blood stage growth of treated (^rapa^) and untreated (^UT^) *ron2*cKO (B) and *ron4*cKO (C) parasites. The graph represents the parasitaemia (mean +/- SEM) in groups of 5 mice. Rapamycin was administered at day 2. **, p < 0.01; ****, p < 0.0001 (Two-way ANOVA). **D-F.** Quantification of midgut sporozoites (MG-SPZ, D), haemolymph sporozoites (HL-SPZ, E) or salivary gland sporozoites (SG-SPZ, F) isolated from mosquitoes infected with untreated (^UT^) or rapamycin treated (^rapa^) *ron2*cKO or *ron4*cKO parasites. The graphs show the number of sporozoites per female mosquito (mean +/- SEM). Each dot represents the mean value obtained in independent experiments after dissection of 30-50 mosquitoes (MG, HL) or 50-70 mosquitoes (SG), respectively. Ns, non-significant; *, p < 0.05; **, p < 0.01 (Two-tailed ratio paired t test). **G-I**. Quantification of excised (mCherry^+^/GFP^−^, red) and non-excised (mCherry^+^/GFP^+^, green) midgut sporozoites (MG-SPZ, G), haemolymph sporozoites (HL-SPZ, H) or salivary gland sporozoites (SG-SPZ, I) isolated from mosquitoes infected with untreated (^UT^) or rapamycin treated (^rapa^) *ron2*cKO and *ron4*cKO parasites. **J.** Quantification of EEFs development *in vitro,* done by microscopy analysis of HepG2 cells infected with sporozoites isolated from either untreated or rapamycin-treated *ron2*cKO and *ron4cKO* infected mosquitoes. The data for rapamycin-treated parasites are represented as percentage of the respective untreated parasites (mean +/- SEM). Each data point is the mean of five technical replicates in one experiment. Ns, non-significant; *, p < 0.05 (Two-tailed ratio paired t test). **K.** Quantification of sporozoite cell traversal activity (% of dextran-positive cells) in untreated and rapamycin-treated *ron2*cKO and *ron4*cKO parasites. The data for rapamycin-treated parasites are represented as percentage of the respective untreated parasites (mean +/- SEM). Each data point is the mean of five technical replicates from one experiment.

We then transmitted *ron2*cKO and *ron4*cKO parasites to mosquitoes, before and after rapamycin treatment respectively. Both parasite lines could colonise the midgut of mosquitoes as evidenced by microscopy imaging of midgut oocysts (**Figure 4-Supplement 3**). Rapamycin treatment of *ron2*cKO and *ron4*cKO parasites before transmission led to a modest reduction of midgut and haemolymph sporozoite numbers (**Figure 4D** and **4E**). However, there was no difference in the percentage of mosquitoes displaying mCherry-labelled pericardial cells (**Figure 4-Supplement 4**), indicating no defect in egress from oocysts for both *ron2*cKO^rapa^ and *ron4*cKO^rapa^ sporozoites. In contrast, the numbers of salivary gland sporozoites were severely reduced with both *ron2*cKO^rapa^ and *ron4*cKO^rapa^ parasites (**Figure 4F**), as observed with *ama1*cKO^rapa^ parasites. As expected, rapamycin treatment before transmission induced robust gene excision in both *ron2*cKO^rapa^ and *ron4*cKO^rapa^ sporozoites (**Figure 4G-I**). Despite reduced invasion we could recover sufficient numbers of *ron2*cKO^rapa^ and *ron4*cKO^rapa^ salivary gland sporozoites to assess host cell invasion *in vitro*. As observed with *ama1*cKO^rapa^ parasites, rapamycin-induced gene excision of *ron2* and *ron4* impaired invasion of HepG2 cells, as shown by reduced EEF numbers (**Figure 4J**). As observed for AMA1-deficient sporozoites, cell traversal activity was preserved in *ron2*cKO^rapa^ and *ron4*cKO^rapa^ sporozoites (**Figure 4K**). Overall, our data support an active role for RON2 and RON4 in invasion of both mosquito salivary glands and hepatocytes, similar to AMA1.

### AMA1-deficient sporozoites show no defect in transcellular migration inside infected salivary glands

In order to get more insights into the colonization of the mosquito salivary glands by sporozoites, we used serial block face-scanning electron microscopy (SBF-SEM) for three-dimensional volume imaging of whole salivary glands isolated from mosquitoes infected with WT (PbGFP) or *ama1*cKO^rapa^ parasites. SBF-SEM data confirmed the lower parasite density in glands infected with *ama1*cKO^rapa^ as compared to WT (**Figure 5A-B**). WT sporozoites were observed inside acinar cells and in the apical secretory cavities, where they clustered in bundles (**Figure 5A** and **Movie 1**). Despite reduced numbers of sporozoites, we observed a similar distribution of *ama1*cKO^rapa^ parasites inside the salivary glands, with both intracellular and intraluminal sporozoites (**Figure 5B** and **Movie 2**). Most of the sporozoites were found lying in direct contact with the cytosol inside acinar cells, without any visible vacuolar membrane (**Figure 5A**). Nevertheless, we also observed some sporozoites surrounded by membranes (**Figure 5C-D**). However, careful examination of the 3D SBF-SEM images revealed that these structures may correspond to invaginations of cellular membranes surrounding portions of intracellular sporozoites, rather than actual vacuoles (**Figure 5C-D** and **Movie 3**). Similar to the WT, *ama1*cKO^rapa^ parasites surrounded by membranes were also found inside acinar cells (**Figure 5E**). We also observed sporozoites present in the secretory cavity and surrounded by a cellular membrane, with both WT (**Figure 5F**) and *ama1*cKO^rapa^ parasites (**Figure 5G**). Altogether these data confirmed the defect of invasion of the mosquito salivary glands by AMA1-deficient sporozoites, but suggested no difference in the distribution of the parasites inside the infected glands or in transcellular migration toward the secretory cavities.

**Figure 5.**
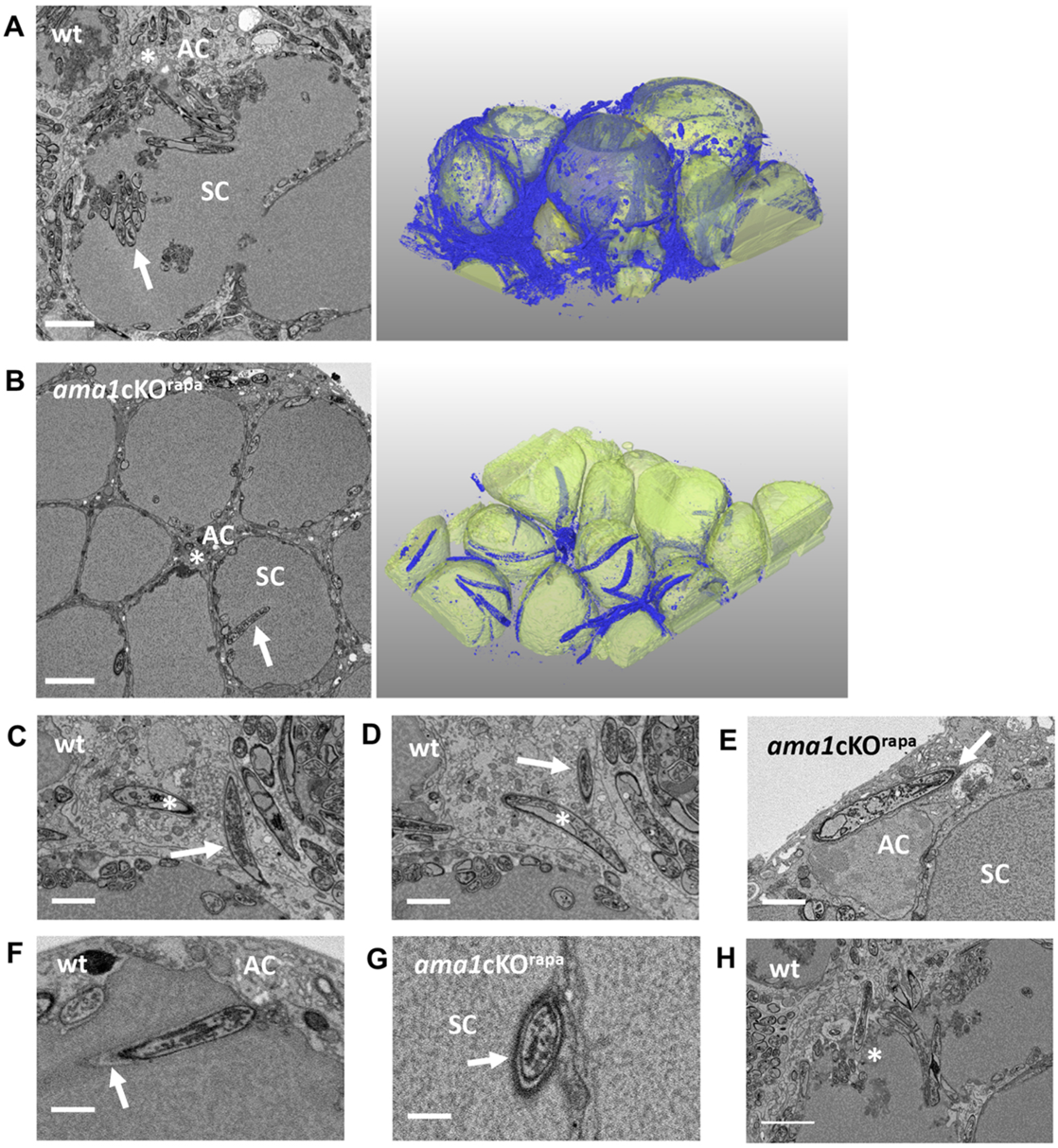
Serial block face-scanning electron microscopy (SBF-SEM) of infected mosquito salivary glands. **A-B**. Representative sections of salivary glands from mosquitoes infected with WT (A) or *ama1*cKO^rapa^ (B) parasites. Scale bars, 5 μm. The volume segmentation shows the secretory cavities (yellow) and sporozoites (blue). WT and AMA1-deficient sporozoites were observed inside the acinar cells (AC, asterisks) and in the secretory cavities (SC, arrows). The volume images corresponding to panel A and panel B are shown in Movie 1 and Movie 2, respectively. **C-D**. SBF-SEM sections from Movie 3, showing WT sporozoites inside salivary gland acinar cells. The first section (C) shows a sporozoite partly surrounded by host cell membranes (arrow), and a second one seemingly contained inside a vacuole (asterisk). The second section (D) shows the same parasites in a different plane, revealing that the second sporozoite is in fact not enclosed in a vacuole but instead is interacting with invaginated host cell membranes (asterisk), while the first parasite now seems surrounded by a membrane (arrow), giving the false impression of being enclosed in a vacuole. Scale bars, 2 μm. **E**. SBF-SEM section showing an intracellular *ama1*cKO^rapa^ sporozoite surrounded by a cellular membrane (arrow). Scale bar, 2 μm. AC, acinar cell; SC, secretory cavity. **F-G**. SBF-SEM sections showing WT (F) and *ama1*cKO^rapa^ (G) sporozoites present inside secretory cavities (SC) and surrounded by cellular membranes (arrows). Scale bars, 1 μm. **H**. SBF-SEM section showing an alteration of the cellular interface with the secretory cavity at the point of entry of multiple WT sporozoites (asterisk). Intraluminal leakage of cytoplasmic material is indicated with an arrow. Scale bar, 5 μm.

### Invasion by AMA1- or RON2-deficient sporozoites is associated with a loss of integrity of the salivary gland epithelium

Interestingly, passage of WT sporozoites from acinar cells to the secretory cavities could be associated with an alteration of the apical cellular membrane integrity, with leakage of cytoplasmic material in the secretory cavity (**Figure 5H**). However, the overall architecture of the infected gland did not seem to be altered despite the presence of numerous WT sporozoites (**Figure 6A**). In contrast, salivary glands from mosquitoes infected with *ama1*cKO^rapa^ parasites showed signs of peripheral cellular damage, with alteration of the basal membrane and cellular vacuolization (**Figure 6B**). To corroborate these observations, we imaged entire salivary glands by fluorescence microscopy after staining of the actin cytoskeleton with phalloidin (**Figure 6C**). Upon examination of salivary glands infected with *ama1*cKO^rapa^ or *ron2*cKO^rapa^, we observed zones where epithelial cells were detached from the basal membrane and retracted, creating pockets suggestive of saliva accumulation, likely as a result of a loss of integrity of the epithelial cell barrier (**Figure 6C**). Such lesions were observed in 9 out of 15 examined salivary gland lobes infected with AMA1- or RON2-deficient sporozoites, but were rarely observed in control parasites (only 1 out of 16 examined salivary glands; P=0.0021, Fisher’s exact test) despite much higher parasite loads (**Figure 6-Supplement 1**). However, heavily infected lobes showed signs of internal remodelling of the actin cytoskeleton (**Figure 6-Supplement 1**), and were prone to rupture during manipulation. Collectively, our data support a role of AMA1 and RONs during invasion of the mosquito salivary glands, which is uncoupled to PV formation, and suggest that sporozoites use the AMA1-RON complex to safely enter the salivary glands without causing cell damage, to ensure successful parasite transmission.

**Figure 6.**
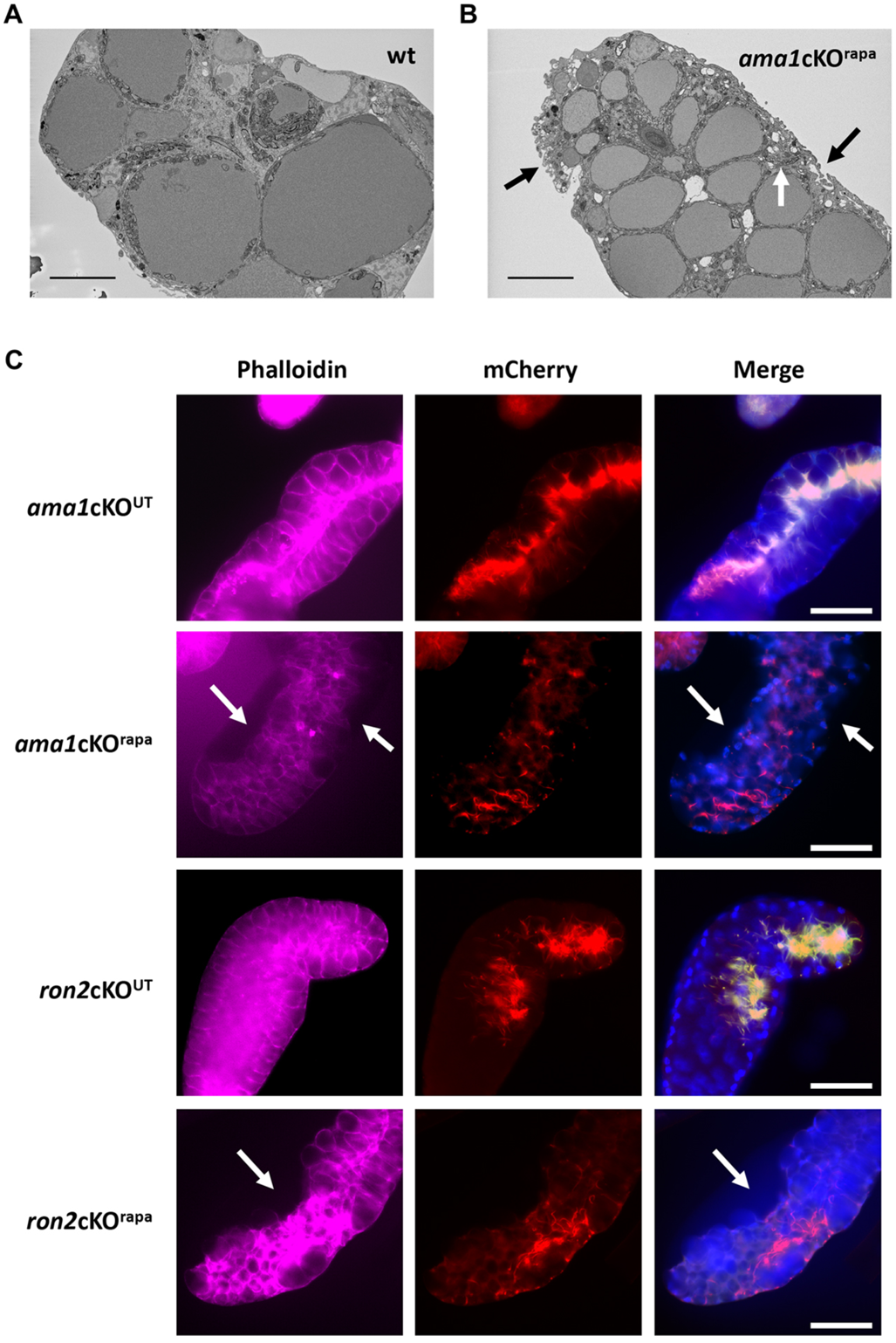
Invasion by AMA1- and RON2-deficient sporozoites is associated with a loss of integrity of the mosquito salivary gland epithelium. **A-B.** SBF-SEM sections of salivary glands infected with WT (I) or *ama1*cKO^rapa^ parasites (J), day 21 post-infection. The gland infected with *ama1*cKO^rapa^ parasites shows signs of cellular damage (black arrows) despite low parasite density. A single intracellular sporozoite is indicated by a white arrow. Scale bars, 10 μm. C. Representative fluorescence microscopy images of salivary gland distal lobes infected with untreated (^UT^) or rapamycin treated (^rapa^) *ama1*cKO or *ron2*cKO parasites, day 16 post-infection. Samples were stained with Phalloidin-iFluor 647 (magenta) and Hoechst 77742 (Blue). The right panels show mCherry (red), GFP (green) and Hoechst (blue) merge images. Zones of retraction of the acinar epithelial cells are visible in the lobes infected with AMA1- and RON2-deficient sporozoites (arrows). Scale bars, 50 μm.

## Discussion

AMA1 and RON proteins play an essential role in *Plasmodium* merozoites during invasion of erythrocytes, where they participate in the formation of the MJ. In contrast, their role in sporozoites was unclear so far. In this study, we exploited the DiCre recombinase system to delete *ama1*, *ron2* or *ron4* genes in *P. berghei* prior to transmission to mosquitoes, allowing subsequent functional investigations in sporozoites. Deletion of the 3’UTR of *ama1* using the DiCre system was not sufficient to abrogate protein expression, a phenomenon that has been observed before with other genes in *P. berghei* and *P. falciparum* (Collins et al., 2013; Ecker et al., 2012), and could explain the discrepancy between our results and previous reports (Giovannini et al., 2011). Therefore, we decided to delete the full-length *ama1* gene in a two-step approach to introduce Lox sites upstream and downstream of the gene of interest in mCherry-expressing PbDiCre parasites, together with a GFP cassette to facilitate monitoring of gene excision. Rapamycin treatment of *ama1*cKO parasites led to a major impairment in blood-stage growth, consistent with an essential role for AMA1 in RBC invasion, but without affecting transmission to mosquitoes. Remarkably, we observed a dramatic reduction in the number of salivary gland sporozoites isolated from *ama1*cKO^rapa^ infected mosquitoes despite comparable midgut and haemolymph sporozoite numbers between *ama1*Con^rapa^ *ama1*cKO^rapa^, showing that AMA1 is important for efficient invasion of the salivary glands, but not egress from oocysts. In addition to a severe defect in salivary gland invasion in the mosquito, AMA1-deficient sporozoites also displayed a defect during invasion of mammalian hepatocytes. Conditional deletion of *ron2* and *ron4* genes resulted in a similar phenotype as with *ama1*, with a severe reduction in salivary gland sporozoite numbers and reduced invasion of hepatocytes. This similar phenotype, combined with mass spectrometry evidence of an interaction between AMA1 and RON proteins, is consistent with AMA1 playing a role together with the RON proteins during sporozoite host cell invasion. It thus appears that the function of AMA1 and RONs cannot be dissociated, unlike previously thought (Giovannini et al., 2011). Our data are in line with those from two studies where a promoter exchange strategy was used to knockdown *ron2*, *ron4* and *ron5* in *P. berghei* sporozoites (Ishino et al., 2019; Nozaki et al., 2020). All three mutants shared a similar phenotype, with a defect in salivary gland invasion and reduced infection of HepG2 cell cultures by haemolymph sporozoites.

Despite poor colonization of the mosquito salivary glands, *ama1*cKO^rapa^ displayed a similar distribution as WT sporozoites inside invaded glands as shown by volume EM imaging, supporting the notion that a functional role of AMA1 (and RONs) is most likely restricted to the initial point of entry into the salivary glands. While we cannot exclude the presence of non-excised *ama1*cKO^rapa^ parasites inside infected glands in the SFB-SEM experiments, these parasites should only represent a minority of salivary gland sporozoites (<10%). Based on our data, we hypothesize that AMA1 is probably secreted from the micronemes before invasion of the glands to form a complex with RON proteins. The similar phenotype observed with *ron2*cKO^rapa^ and *ron4*cKO^rapa^ parasites further indicate that rhoptry secretion also precedes entry into the salivary glands. Previous ultrastructural imaging studies of sporozoites have in fact reported the presence of four or more rhoptries in midgut-derived sporozoites, as opposed to two in mature salivary gland sporozoites (Kudryashev et al., 2010; Schrevel et al., 2008; Sinden and Strong, 1978; Tokunaga et al., 2019). These observations suggest the possibility of a rhoptry discharge event occurring before or during invasion of the glands. However, it is also possible that rhoptries partially secrete their contents, in a kiss-and-run process, in order to facilitate entry into the glands, a process that remains difficult to capture with the current tools available.

Although reminiscent of blood-stage merozoites, where the discharge of RON proteins has been associated with the formation of the MJ (Lamarque et al., 2011; Riglar et al., 2011), it is not known whether sporozoites form a MJ during invasion of mosquito salivary glands. A previous electron microscopy analysis of the salivary glands of *Aedes aegypti* mosquitoes infected with avian *P. gallinaceum* documented sporozoites entering the salivary glands through an invagination of the basal lamina while forming a junctional area between the anterior tip of the sporozoite and the plasma membrane of the acinar cells (Pimenta et al., 1994). In another study, *P. falciparum* sporozoites were observed penetrating salivary glands of *Anopheles stephensi* mosquitoes through holes in the basal membrane without causing any obvious damage to the gland (Meis et al., 1992). Aside from the MJ, there are also conflicting observations on the formation of vacuoles during salivary gland invasion. Pimenta *et al*. reported that newly invaded sporozoites found in the cytoplasm of acinar cells were surrounded by a vacuole while those that had entered the secretory cavity were either devoid of a vacuole or present inside disintegrating vacuoles (Pimenta et al., 1994). Similar vacuoles were occasionally observed surrounding intracellular *P. falciparum* sporozoites in the salivary glands of *A. gambiae* mosquitoes (Sinden and Strong, 1978). Several other studies looking at sporozoites inside the salivary gland secretory cells of infected mosquitoes failed to observe a PV around sporozoites (Posthuma et al., 1989; Sterling et al., 1973; Wells and Andrew, 2019). Here we could resolve in part these discrepancies using volume electron microscopy, which documented the presence of cellular membrane invaginations around intracellular sporozoites, sometimes resulting in the appearance of a vacuole. Our data suggest that during transcellular migration *Plasmodium* sporozoites can push the cell membranes to reach the secretory cavities, a process that appears to be independent of AMA1 and does not involve the formation of a canonical PV.

While our data indicate that inside the acinar cells sporozoites do not reside in a PV, we cannot exclude that haemolymph sporozoites initially enter the salivary glands by forming a transient vacuole. In this regard, mature sporozoites can form transient vacuoles during traversal of mammalian cells (Bindschedler et al., 2020; Risco-Castillo et al., 2015). However, this process is not associated with rhoptry discharge (Risco-Castillo et al., 2015), and we show here that neither AMA1 nor RONs are required for cell traversal activity, suggesting that distinct mechanisms drive invasion of the salivary glands and traversal through mammalian cells. Nevertheless, it is possible that sporozoites use a common mechanism for transcellular migration once inside the mosquito salivary glands and in the mammalian host, both associated with membrane wrapping around traversing parasites ((Risco-Castillo, 2015) and this study).

Interestingly, infection of the mosquito salivary glands with AMA1- or RON2-deficient sporozoites was associated with a loss of integrity of the epithelium. This suggests that during sporozoite entry in the salivary gland, the role of AMA1-RONs could be to maintain a sealed junction around the parasite, to allow invasion without creating a leak, thus preventing cell damage. These observations are reminiscent of the erythrocyte lysis observed during invasion of AMA1-depleted *P. falciparum* merozoites (Collins et al., 2020). Our data thus provide a possible molecular basis to explain how thousands of sporozoites can colonize the salivary glands of a single mosquito without causing overt tissue damage. While sporozoites do not reside in a PV inside the salivary glands, they can remain in the salivary cavities for several days before they are transmitted. Therefore, harmless entry in the glands is likely essential to ensure parasite transmission.

Despite the significant reduction in numbers, a minor proportion of *ama1*cKO^rapa^, *ron2*cKO^rapa^ and *ron4*cKO^rapa^ sporozoites could still invade the salivary glands of infected mosquitoes, suggesting that these sporozoites somehow adapted to the loss of AMA1, RON2 or RON4, perhaps through the use of alternative adhesion or invasion ligands. One can assume that the residual invasion we observed is the outcome of some degree of plasticity in AMA-RON interactions, whereby paralogs may compensate for the function of AMA1 and/or RONs, as observed in *T. gondii* (Lamarque et al., 2014). While there is no known paralog of RON2 in *Plasmodium*, the Membrane Associated Erythrocyte Binding-Like protein (MAEBL) contains two AMA1-like domains (Kappe et al., 1998), and was in fact reported to be essential for invasion of the salivary glands (Kariu et al., 2002; Saenz et al., 2008). Interestingly, MAEBL was not identified by co-immunoprecipitation in the RON2, RON4, RON5 complex in oocyst derived (Nozaki et al., 2020) or salivary gland (this study) sporozoites, and *ama1*cKO^rapa^ sporozoites fail to invade the mosquito salivary glands, thus arguing against a compensatory role for MAEBL in AMA1-deficient sporozoites. Yet, we cannot exclude that MAEBL and AMA1-RONs function at the same step and cooperate for successful invasion of the salivary gland and possibly hepatocytes. Alternatively, MAEBL may play a role upstream of AMA1 and RONs, for example during sporozoite attachment to the salivary glands, together with other proteins such as TRAP (Ghosh et al., 2009), triggering subsequent rhoptry secretion for AMA1-RON-dependent invasion of the gland.

When tested on hepatocyte cell cultures, only a minor proportion of *ama1*cKO^rapa^, *ron2*cKO^rapa^ and *ron4*cKO^rapa^ salivary gland sporozoites productively invaded and developed into EEFs, suggesting that the functional role of AMA1, RON2 and RON4 during invasion is conserved across invasive stages. The defect in hepatocyte invasion however was less pronounced in comparison to that observed for the salivary glands, implying either a possible adaptation of these parasites, or selection of the most invasive sporozoites (including the small number of non-excised parasites), or simply a differential dependency on AMA1-RONs during liver infection, which could relate to different membrane properties. Consistent with our results, a previous study has shown that anti-AMA1 only partially inhibited *P. falciparum* infection of human hepatocytes *in vitro* (Silvie et al., 2004). Interestingly, knockdown of RON2 in sporozoites was shown to affect cell traversal and hepatocyte invasion, both *in vitro* and *in vivo*, with the authors implying that loss of RON2 affected attachment to both the salivary glands and hepatocytes, thereby influencing invasion (Ishino et al., 2019). An earlier report on *P. falciparum* sporozoites showed that interfering with the AMA1-RON2 interaction affected host cell traversal (Yang et al., 2017). However, in our study, *ama1*cKO^rapa^, *ron2*cKO^rapa^ and *ron4*cKO^rapa^ sporozoites showed no defect in cell traversal but were impaired in productive invasion. While these differences in phenotypes could be attributed to differences between *P. falciparum* and *P. berghei*, it is possible that the use of salivary gland sporozoites in our study versus those obtained from the haemolymph by Ishino *et al*. accounted for the difference in observations for cell traversal between experiments.

Based on our findings, we propose a model where *Plasmodium* sporozoites use the AMA1-RON complex twice, in the mosquito and mammalian hosts. First, AMA1 and RONs could mediate the safe entry of sporozoites into the salivary glands, possibly via the formation of a transient vacuole, in a cell-specific manner and without compromising the cell membrane integrity, to ensure successful colonization of the glands and subsequent parasite transmission. This model fits with previous reports showing that sporozoites can massively infect salivary glands without causing cellular damage (Posthuma et al., 1989; Wells and Andrew, 2019). This crossing event would differ from the cell traversal activity of mature sporozoites in the mammalian host, which is associated with a loss of membrane integrity and cell death (Formaglio et al., 2014). Following sporozoite inoculation into the mammalian host, AMA1 and RONs facilitate productive invasion of hepatocytes, likely through the formation of a canonical MJ that leads to the formation of the PV where the parasite can replicate into merozoites. Colonization of the salivary glands and productive invasion of hepatocytes are distinct mechanisms, involving transcellular migration versus establishment of a replicative vacuole, respectively. However, both events likely require tight membrane sealing around the invading parasite, a function that could rely on the AMA1-RON complex. Our study reveals that the contribution of AMA1 and RON proteins is conserved across *Plasmodium* invasive stages. Pre-clinical studies have shown that vaccination with the AMA1-RON2 complex induces functional antibodies that better recognize AMA1 as it appears complexed with RON2 during merozoite invasion, providing an attractive vaccine strategy against *Plasmodium* blood stages (Srinivasan et al., 2017, 2014). Our results indicate that the AMA1-RON complex might also be considered as a potential target to block malaria transmission.

## Materials and methods

### Mice

Female Swiss mice (6–8 weeks old, from Janvier Labs) were used for all routine parasite infections. All animal work was conducted in strict accordance with the Directive 2010/63/EU of the European Parliament and Council ‘On the protection of animals used for scientific purposes’. Protocols were approved by the Ethical Committee Charles Darwin N°005 (approval #7475-2016110315516522).

### Parasites

Conditional genome editing was performed in the *P. berghei* (ANKA strain) PbDiCre line, obtained after integration of mCherry and DiCre expression cassettes at the dispensable *p230p* locus (Fernandes et al., 2020). Two additional lines expressing RON4-mCherry (bioRxiv 2021.10.25.465731) and/or GFP (Manzoni et al., 2014) were used for immunoprecipitation and electron microscopy experiments, respectively. Parasites were maintained in mice through intraperitoneal injections of frozen parasite stocks. *Anopheles stephensi* mosquitoes were reared at 24°C with 80 % humidity and permitted to feed on infected mice that were anaesthetised, using standard methods of mosquito infection as previously described (Ramakrishnan et al., 2013). Post feeding, *P. berghei*-infected mosquitoes were kept at 21°C and fed daily on a 10% sucrose solution.

### Host cell cultures

HepG2 cells (ATCC HB-8065) were cultured in DMEM supplemented with 10% foetal calf serum, 1% Penicillin-Streptomycin and 1% L-Glutamine as previously described (Silvie et al., 2007), in culture dishes coated with rat tail collagen I (Becton-Dickinson).

### Vector construction

In order to target different genes of interest, we first generated a generic plasmid, pDownstream1Lox (Addgene #164574), containing a GFP-2A-hDHFR cassette under the control of a *P. yoelii hsp70* promoter and followed by the 3’UTR of *P. berghei calmodulin (cam)* gene and a single LoxN site. The plasmid also contains a yFCU cassette to enable the elimination of parasites carrying episomes by negative selection with 5-fluorocytosine.

The *ama1*Con plasmid was designed to excise only ∼30 bp downstream of *P. berghei ama1* 3’UTR. Two fragments were inserted on each side of the GFP-2A-hDHFR cassette of the pDownstream1Lox plasmid: a 5’ homology region (HR) homologous to the terminal portion of *ama1* (ORF and 3’ UTR) followed by a single LoxN site, and a 3’ HR homologous to a sequence downstream of the 3’ UTR of *ama1* gene. The *ama1*Δutr plasmid was assembled similarly to the *ama1*Con construct except that the 5’ HR consisted in the terminal portion of *ama1* ORF followed by a LoxN site and the 3’ UTR of *P. yoelii ama1*, to allow excision of the 3’UTR upon rapamycin activation of DiCre. The *ama1*cKO plasmid was designed to introduce a single LoxN site upstream of *ama1* in the *ama1*Con^rapa^ parasites, which already contained a residual LoxN site downstream of the gene. To generate the *ama1*cKO plasmid, the pDownstream1Lox vector was first modified to remove the downstream LoxN site. Then, a 5’ HR and a 3’ HR, both homologous to sequences located upstream of *ama1* gene, were cloned into the modified plasmid on each side of the GFP-2A-hDHFR, with a single LoxN site introduced upstream of the GFP-2A-hDHFR cassette.

To generate *ron2*cKO and *ron4*cKO constructs, two separate plasmids, P1 and P2, were generated to insert a LoxN site upstream of the promoter and downstream of the gene of interest, respectively, in two consecutive transfections. P1 plasmids were constructed by insertion of 5’ and 3’ HR on each side of the GFP-2A-hDHFR cassette in the pDownstream1Lox plasmid, with a second LoxN site introduced upstream of the GFP cassette. The 5’ HR and 3’ HR correspond to consecutive fragments located in the promoter region of the GOI. Because the intergenic sequence between *ron4* gene and its upstream gene is short, and in order to maintain expression of the upstream gene and exclude any unwanted duplication and spontaneous recombination events, we introduced the 5’ HR of *ron4* in two fragments, with fragment 1 corresponding to the region just upstream of the ORF while fragment 2 corresponded to the 3’ UTR from the *P. yoelii* ortholog of the upstream gene. P2 plasmids were constructed in a similar manner by insertion of a 5’ HR and a 3’HR on each side of the GFP-2A-hDHFR cassette in the pDownstream1Lox plasmid. The 3’ HR regions corresponded to the 3’ UTR sequences of *RON2* or *RON4*, respectively. For both target genes, the 5’ HR was divided into two fragments, where fragment 1 corresponded to the end of the ORF followed by a triple Flag tag, and fragment 2 corresponded to the 3’ UTR from the *P. yoelii* ortholog gene, in order to avoid duplication of the 3’ UTR region and spontaneous recombination.

All plasmid inserts were amplified by PCR using standard PCR conditions and the CloneAmp HiFi PCR premix (Takara). Following a PCR purification step (QIAquick PCR purification kit), the fragments were sequentially ligated into the target vector using the In-Fusion HD Cloning Kit (Clontech). The resulting plasmid sequences were verified by Sanger sequencing (GATC Biotech) and linearised before transfection. All the primers used for plasmid assembly are listed in **Table S2**.

### Parasite transfection

For parasite transfection, schizonts purified from an overnight culture of PbDiCre parasites were transfected with 5–10 µg of linearised plasmid by electroporation using the AMAXA Nucleofector device (Lonza, program U033), as previously described (Janse et al., 2006), and immediately injected intravenously into the tail vein of Swiss mice. For selection of resistant transgenic parasites, pyrimethamine (35 mg/L) and 5-flurocytosine (0.5 mg/ml) were added to the drinking water and administered to mice, one day after transfection. Transfected parasites were sorted by flow cytometry on a FACSAria II (Becton-Dickinson), as described (Manzoni et al., 2014), and cloned by limiting dilutions and injections into mice. The parasitaemia was monitored daily by flow cytometry and the mice sacrificed at a parasitaemia of 2-3%. The mice were bled and the infected blood collected for preparation of frozen stocks (1:1 ratio of fresh blood mixed with 10% Glycerol in Alsever’s solution) and isolation of parasites for genomic DNA extraction, using the DNA Easy Blood and Tissue Kit (Qiagen), according to the manufacturer’s instructions. Specific PCR primers were designed to check for wild-type and recombined loci and are listed in **Table S2**. Genotyping PCR reactions were carried out using Recombinant Taq DNA Polymerase (5U/µl from Thermo Scientific) and standard PCR cycling conditions.

### *In vivo* analysis of conditional mutants

DiCre recombinase mediated excision of targeted DNA sequences *in vivo* was achieved by a single oral administration of 200µg rapamycin (1mg/ml stock, Rapamune, Pfizer) to mice. Excision of the GFP cassette in blood stage parasites was monitored by flow cytometry using a Guava EasyCyte 6/2L bench cytometer equipped with 488 nm and 532 nm lasers (Millipore) to detect GFP and mCherry, respectively. To analyse parasite development in the mosquito, rapamycin was administered to infected mice 24 hours prior to transmission to mosquitoes, as described (Fernandes et al., 2020). Midguts were dissected out at day 14 post infection. The haemolymph was collected by flushing the haemocoel with complete DMEM, day 14 to 16 post infection. Salivary gland sporozoites were collected between 21–28 days post feeding from infected mosquitoes, by hand dissection and homogenisation of isolated salivary glands in complete DMEM. Live samples (infected mosquito midguts or salivary glands, sporozoites) were mounted in PBS and visualised live using a Zeiss Axio Observer.Z1 fluorescence microscope equipped with a LD Plan-Neofluar 403/0.6 Corr Ph2 M27 objective. The exposure time was set according to the positive control and maintained for both untreated and rapamycin-treated parasites, in order to allow comparisons. All images were processed with ImageJ for adjustment of contrast.

### *In vitro* sporozoite assays

HepG2 cells were seeded at a density of 30,000 cells/well in a 96-well plate for flow cytometry analysis or 10,000 cells/well in 8 well µ-slide (IBIDI) for immunofluorescence assays, 24 hours prior to infection with sporozoites. On the day of the infection, the culture medium in the wells was discarded and fresh complete DMEM was added along with 10,000 sporozoites, followed by incubation for 3 hours at 37°C. After 3 hours, the wells were washed twice with complete DMEM and then incubated for another 24-48 hours at 37°C and 5% CO_2_. For quantification of EEF numbers, the cells were trypsinised after two washes with PBS, followed by addition of complete DMEM and one round of centrifugation at 4°C. After discarding the supernatant, the cells were either directly re-suspended in complete DMEM for flow cytometry, or fixed with 2% PFA for 10 minutes, subsequently washed once with PBS and then re-suspended in PBS for FACS acquisition. For quantification of traversal events, fluorescein-conjugated dextran (0.5mg/ml, Life Technologies) was added to the wells along with sporozoites followed by an incubation at 37°C for 3 hours. After 3 hours, the cells were washed twice with PBS, trypsinised and resuspended in complete DMEM for analysis by flow cytometry.

### RON4 immunoprecipitation and mass spectrometry

Freshly dissected RON4-mCherry sporozoites were lysed on ice for 30 min in a lysis buffer containing 0.5% w/v NP40 and protease inhibitors. After centrifugation (15,000 × g, 15 min, 4°C), supernatants were incubated with protein G-conjugated sepharose for preclearing overnight. Precleared lysates were subjected to mCherry immunoprecipitation using RFP-Trap beads (Chromoteck) for 2h at 4°C, according to the manufacturer’s protocol. PbGFP parasites with untagged RON4 were used as a control. After washes, proteins on beads were eluted in 2X Laemmli and denatured (95°C, 5min). After centrifugation, supernatants were collected for further analysis. Samples were subjected to a short SDS-PAGE migration, and gel pieces were processed for protein trypsin digestion by the DigestProMSi robot (Intavis), as described (Hamada et al., 2021). Peptide were separated on an Aurora UHPLC column from IonOpticks (25 cm x 75 μm, C18), using a 30 min gradient from 3 to 32% ACN with 0.1% formic acid, and analysed on a timsTOF PRO mass spectrometer (Bruker). Mascot generic files were processed with X!Tandem pipeline (version 0.2.36) using the PlasmoDB_PB_39_PbergheiANKA database, as described (Hamada et al., 2021).

### Immunofluorescence assays

Blood-stage schizonts were fixed with 4% PFA and 0.0075% glutaraldehyde for 30 mins at 37°C with constant shaking. The samples were then quenched/permeabilised with 125mM glycine /0.1% Triton X-100 for 15 minutes, blocked with PBS/3% BSA, then incubated with Rat anti-AMA1 antibodies (1:250, clone 28G2, MRA-897A, Bei Resources) followed by Alexa Fluor goat anti-rat 405 antibodies (1:1000, Life Technologies). The samples were mounted in PBS and immediately visualised under a fluorescence microscope. Sporozoites were resuspended in PBS, added on top of poly-L-lysine coated coverslips and allowed to air dry. The sporozoites were then fixed with 4% PFA for 30 mins, followed by quenching with 0.1M glycine for 30 mins and two washes with PBS. In the next step, the sporozoites were permeabilised with 1% Triton-X for 5 mins, washed twice with PBS, then blocked with PBS 3%BSA for 1hr at RT and incubated with anti-AMA1 antibody (1:250) diluted in blocking solution. Following 3 washes with PBS, the sporozoites were incubated with the secondary antibody (anti-Rat Alexa Fluor 647) diluted in blocking solution. Following 3 washes with PBS, the coverslips were mounted onto a drop of prolong diamond anti-fade mounting solution (Life Technologies), sealed with nail polish and imaged using a fluorescence microscope. Infected HepG2 cell cultures were washed twice with PBS, then fixed with 4% PFA for 20 minutes, followed by two washings with PBS and incubation with goat anti-UIS4 primary antibody (1:500, Sicgen), followed by donkey anti-goat Alexa Fluor 594 secondary antibody (1:1000, Life Technologies). For fluorescence imaging of entire glands, freshly dissected salivary glands were fixed in 4% PFA for 30 minutes and permeabilized in acetone for 90 seconds, as described (Wells and Andrew, 2019). Samples were incubated with Phalloidin-iFluor 647 (Abcam) and Hoechst 77742 (Life Technologies) overnight at 4°C, washed in PBS and mounted in glycerol before imaging. Acquisitions were made on a Zeiss Axio Observer Z1 fluorescence microscope using the Zen software (Zeiss). Images were processed with ImageJ for adjustment of contrast.

### Serial block face-scanning electron microscopy

For Serial Block Face-Scanning Electron Microscopy (SBF-SEM), salivary glands were isolated from mosquitoes infected with PbGFP or *ama1*cKO^rapa^ parasites, and fixed in 0.1 M cacodylate buffer containing 3% PFA and 1% glutaraldehyde during 1 hour at room temperature. Intact salivary glands were then prepared for SBF-SEM (NCMIR protocol) (Deerinck et al., 2010) as follows: samples were post-fixed for 1 hour in a reduced osmium solution containing 1% osmium tetroxide, 1.5% potassium ferrocyanide in PBS, followed by incubation with a 1% thiocarbohydrazide in water for 20 minutes. Subsequently, samples were stained with 2% OsO4 in water for 30 minutes, followed by 1% aqueous uranyl acetate at 4 °C overnight. Samples were then subjected to en bloc Walton’s lead aspartate staining (Walton, 1979), and placed in a 60 °C oven for 30 minutes. Samples were then dehydrated in graded concentrations of ethanol for 10 minutes in each step. The samples were infiltrated with 30% agar low viscosity resin (Agar Scientific Ltd, UK) in ethanol, for 1 hour, 50% resin for 2 hours and 100% resin overnight. The resin was then changed and the samples were further incubated during 3 hours, prior to inclusion by flat embedding between two slides of Aclar® and polymerization for 18 hours at 60 °C. The polymerized blocks were mounted onto aluminum stubs for SBF-SEM imaging (FEI Microtome 8 mm SEM Stub, Agar Scientific), with two-part conduction silver epoxy kit (EMS, 12642-14). For imaging, samples on aluminum stubs were trimmed using an ultramicrotome and inserted into a TeneoVS SEM (ThermoFisher Scientific). Acquisitions were performed with a beam energy of 2 kV, 400 pA current, in LowVac mode at 40 Pa, a dwell time of 1 µs per pixel at 10 nm pixel size. Sections of 50 nm were serially cut between images. Data acquired by SBF-SEM were processed using Fiji and Amira (ThermoFisher Scientific). Data alignment and manual segmentation were performed using Amira.

### Quantification and statistical analysis

*In vitro* experiments were performed with a minimum of three technical replicates per experiment. Statistical significance was assessed by two-way ANOVA, one-way ANOVA followed by Tukey’s multiple comparisons, or ratio paired t tests, as indicated in the figure legends. All statistical tests were computed with GraphPad Prism 5 (GraphPad Software). The quantitative data used to generate the figures and the statistical analysis are presented in **Table S3.**

## Acknowledgements

We thank Jean-François Franetich, Maurel Tefit and Thierry Houpert for rearing of mosquitoes, and Maryse Lebrun for helpful discussions. The following reagent was obtained through BEI Resources, NIAID, NIH: Monoclonal Anti-*Plasmodium* Apical Membrane Antigen 1, Clone 28G2 (produced *in vitro*), MRA-897A, contributed by Alan W. Thomas. This work was funded by grants from the Laboratoire d’Excellence ParaFrap (ANR-11-LABX-0024), the Agence Nationale de la Recherche (ANR-16-CE15-0004 and ANR-16-CE15-0010) and the Fondation pour la Recherche Médicale (EQU201903007823). The authors acknowledge the Conseil Régional d’Ile-de-France, Sorbonne Université, the National Institute for Health and Medical Research (INSERM) and the Biology, Health and Agronomy Infrastructure (IBiSA) for funding the timsTOF PRO. We acknowledge the ImagoSeine core facility of the Institut Jacques Monod, member of the France BioImaging infrastructure (ANR-10-INBS-04) and GIS-IBiSA, and funded by Région Ile-de-France (TeneoVS). ML was supported by a ‘DIM 1Health’ doctoral fellowship awarded by the Conseil Régional d’Ile-de-France. AW is supported by the ATIP-Avenir program.

## Supplemental material

**Figure 1 – Supplement 1.**
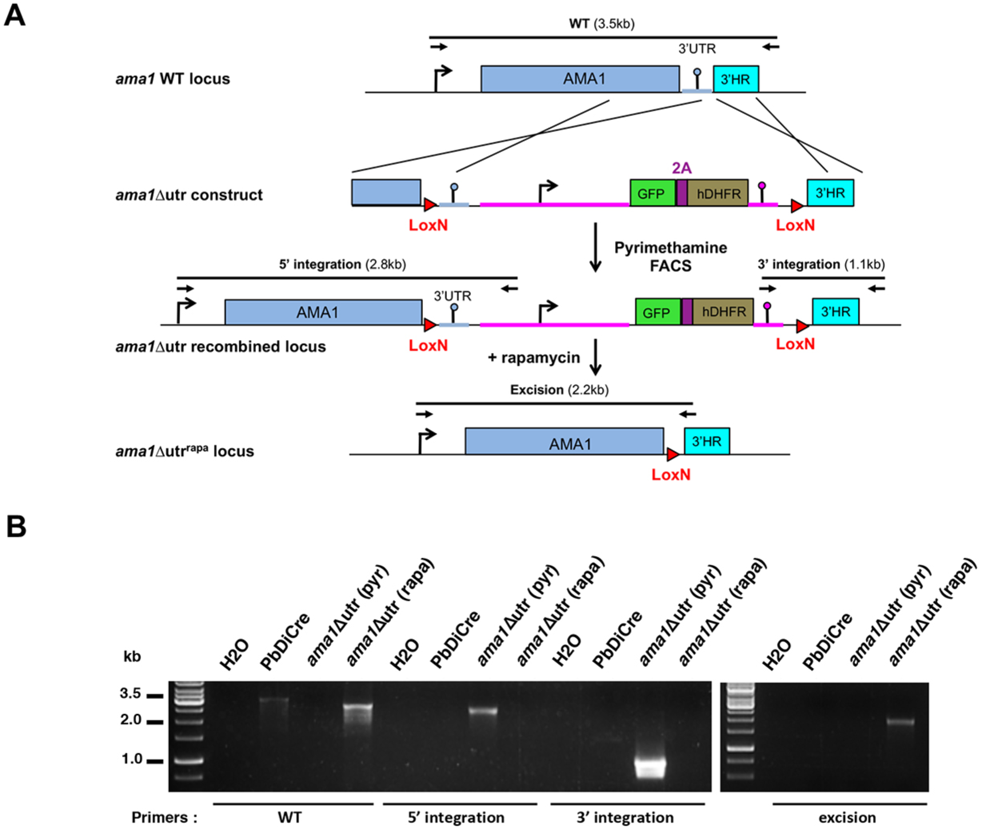
Generation of *ama1*Δutr parasites using the DiCre system. **A.** Strategy to generate *ama1*Δutr parasites. The wild-type locus of *P. berghei AMA1* in the PbDiCre parasite line was targeted with a *ama1*Δutr replacement plasmid containing 2 Lox sites and 5’ and 3’ homologous sequences inserted on each side of a GFP-2A-hDHFR cassette. Upon double crossover recombination, the LoxN sites are inserted upstream of the 3’ UTR and downstream of the GFP-2A-hDHFR cassette, respectively. Activation of the DiCre recombinase with rapamycin results in excision of the 3’ UTR together with the GFP-2A-hDHFR cassette. Genotyping primers and expected PCR fragments are indicated by arrows and lines, respectively. **B.** Genotyping of parental PbDiCre and *ama1*Δutr transfected parasites after pyrimethamine selection (pyr) and after rapamycin treatment (rapa) of the final population. Parasite genomic DNA was analysed by PCR using primer combinations specific for the unmodified locus (WT), the 5’ integration, 3’ integration or excision events.

**Figure 1 – Supplement 2.**
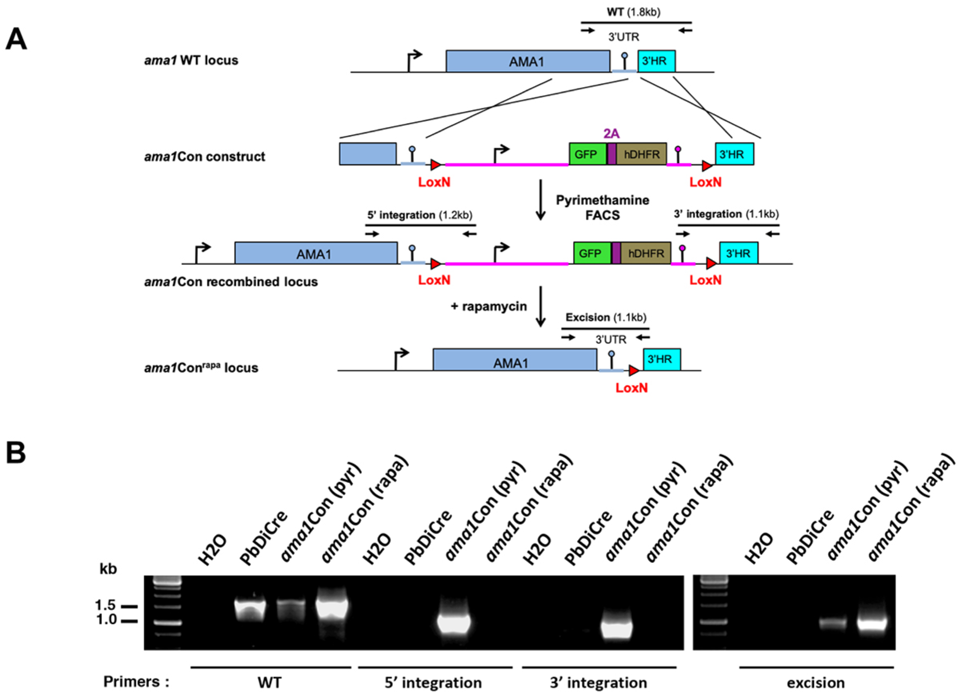
Generation of *ama1*Con parasites using the DiCre system. **A.** Strategy to generate *ama1*Con parasites. The construct is similar to the *ama1*Δutr construct, except that the first LoxN site is located downstream of the 3’ UTR. Upon rapamycin-induced excision, the *AMA1* locus remains intact. **B.** Genotyping of parental PbDiCre and *ama1*Con transfected parasites after pyrimethamine selection (pyr) and after rapamycin treatment (rapa) of the final population. Parasite genomic DNA was analysed by PCR using primer combinations specific for the unmodified locus (WT), the 5’ integration, 3’ integration or excision events.

**Figure 1 – Supplement 3.**
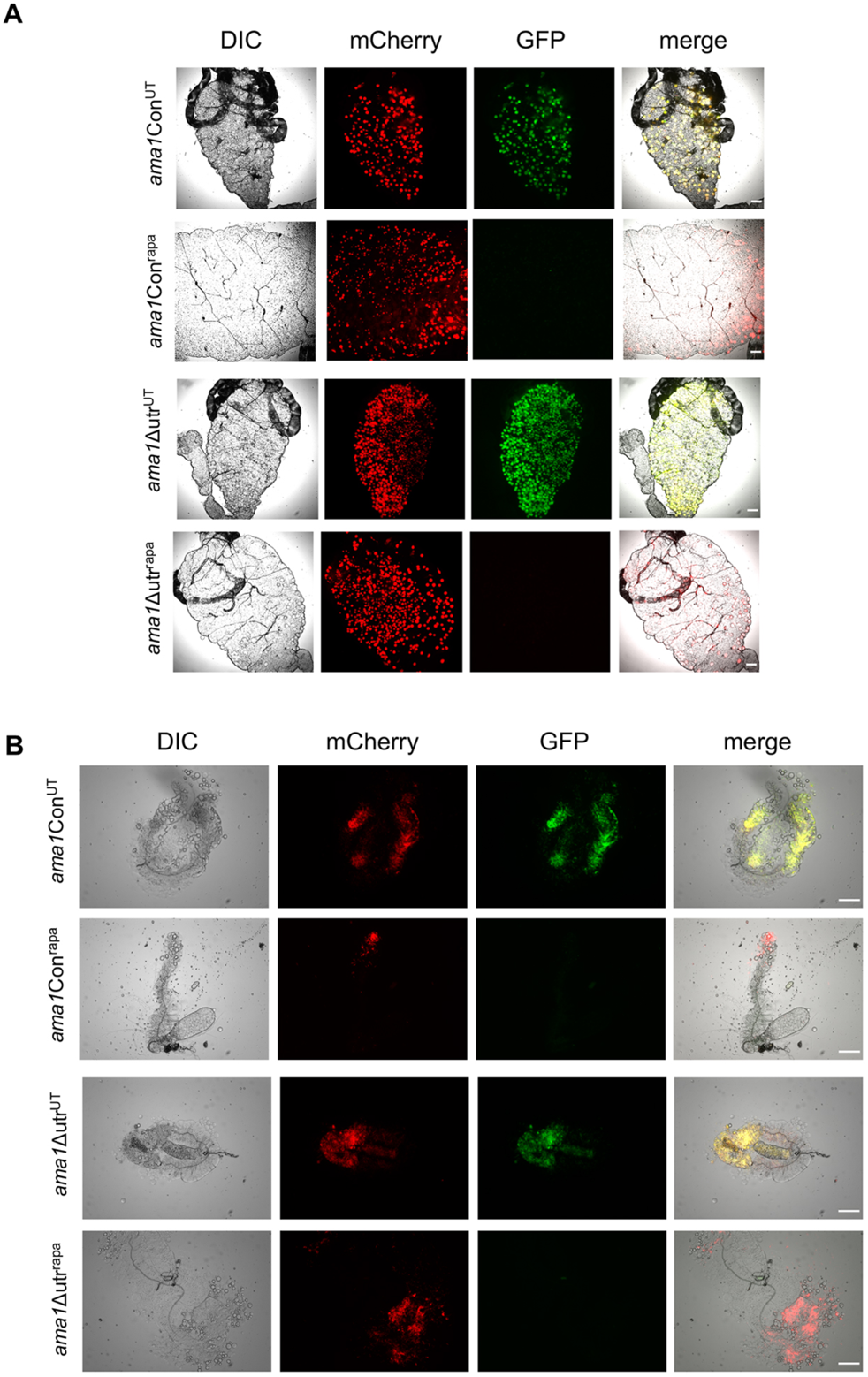
Imaging of *ama1*Con and *ama1*Δutr mosquito stages. **A.** Fluorescence microscopy images of midguts from mosquitoes infected with untreated (UT) or rapamycin-treated (rapa) *ama1*Con and *ama1*Δutr parasites. Scale bar = 200 µm. **B.** Fluorescence microscopy images of salivary glands from mosquitoes infected with untreated (UT) or rapamycin-treated (rapa) *ama1*Con and *ama1*Δutr parasites. Scale bar = 200 µm.

**Figure 2 – Supplement 1.**
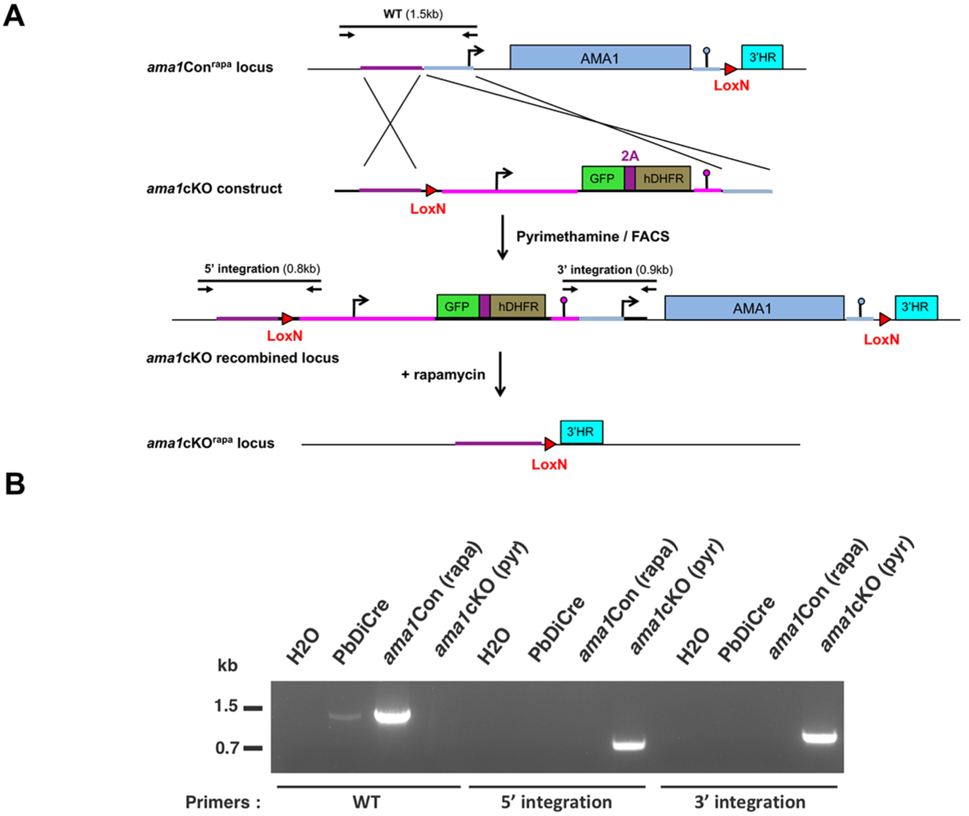
Generation of *ama1*cKO parasites using the DiCre system. **A.** Strategy to generate *ama1*cKO parasites. The *AMA1* locus in *ama1*Con^rapa^ parasites was targeted with a *ama1*cKO replacement plasmid containing a single LoxN site and 5’ and 3’ homologous sequences inserted on each side of a GFP-2A-hDHFR cassette. Upon double crossover recombination, a second LoxN site is inserted upstream of the GFP-2A-hDHFR cassette and *AMA1* gene. Activation of the DiCre recombinase with rapamycin results in excision of the entire *AMA1* gene together with the GFP-2A-hDHFR cassette. Genotyping primers and expected PCR fragments are indicated by arrows and lines, respectively. **B.** Genotyping of PbDiCre, *ama1*Con^rapa^ (parental) and *ama1*cKO parasites. Parasite genomic DNA was analysed by PCR using primer combinations specific for the unmodified locus (WT), the 5’ integration and 3’ integration events.

**Figure 2 – Supplement 2.**
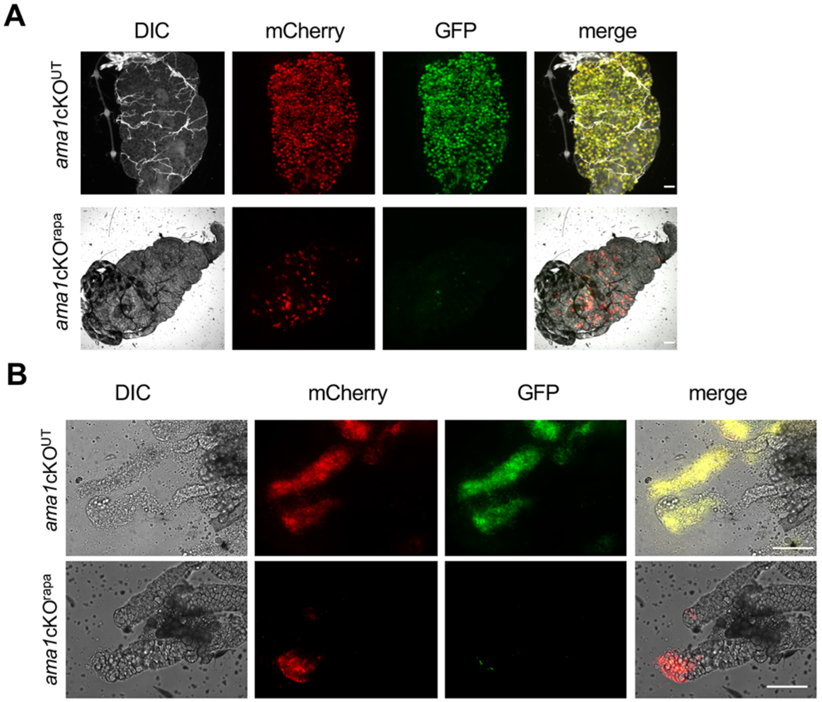
Imaging of *ama1*cKO mosquito stages. **A.** Fluorescence microscopy of midguts from mosquitoes infected with untreated (UT) or rapamycin-treated (rapa) *ama1*cKO parasites. Scale bar = 200 µm. **B.** Fluorescence microscopy of salivary glands isolated from mosquitoes infected with untreated (UT) or rapamycin-treated (rapa) *ama1*cKO parasites. Scale bar = 200 µm.

**Figure 2 – Supplement 3.**
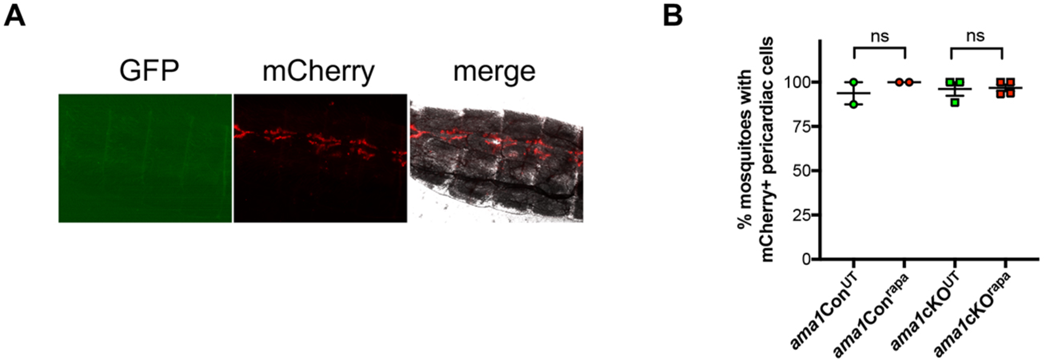
Analysis of mosquito pericardial structures. **A.** Imaging of the abdomen of a mosquito infected with rapamycin treated *ama1*cKO parasites, after removal of the midgut, showing mCherry-labelled pericardial structures. **B.** Quantification of mosquitoes with mCherry-labelled pericardial cells at D21 post-infection with untreated (UT) or rapamycin-treated (rapa) *ama1*Con and *ama1*cKO parasites. Ns, non-significant (Two-tailed ratio paired t test).

**Figure 4 – Supplement 1.**
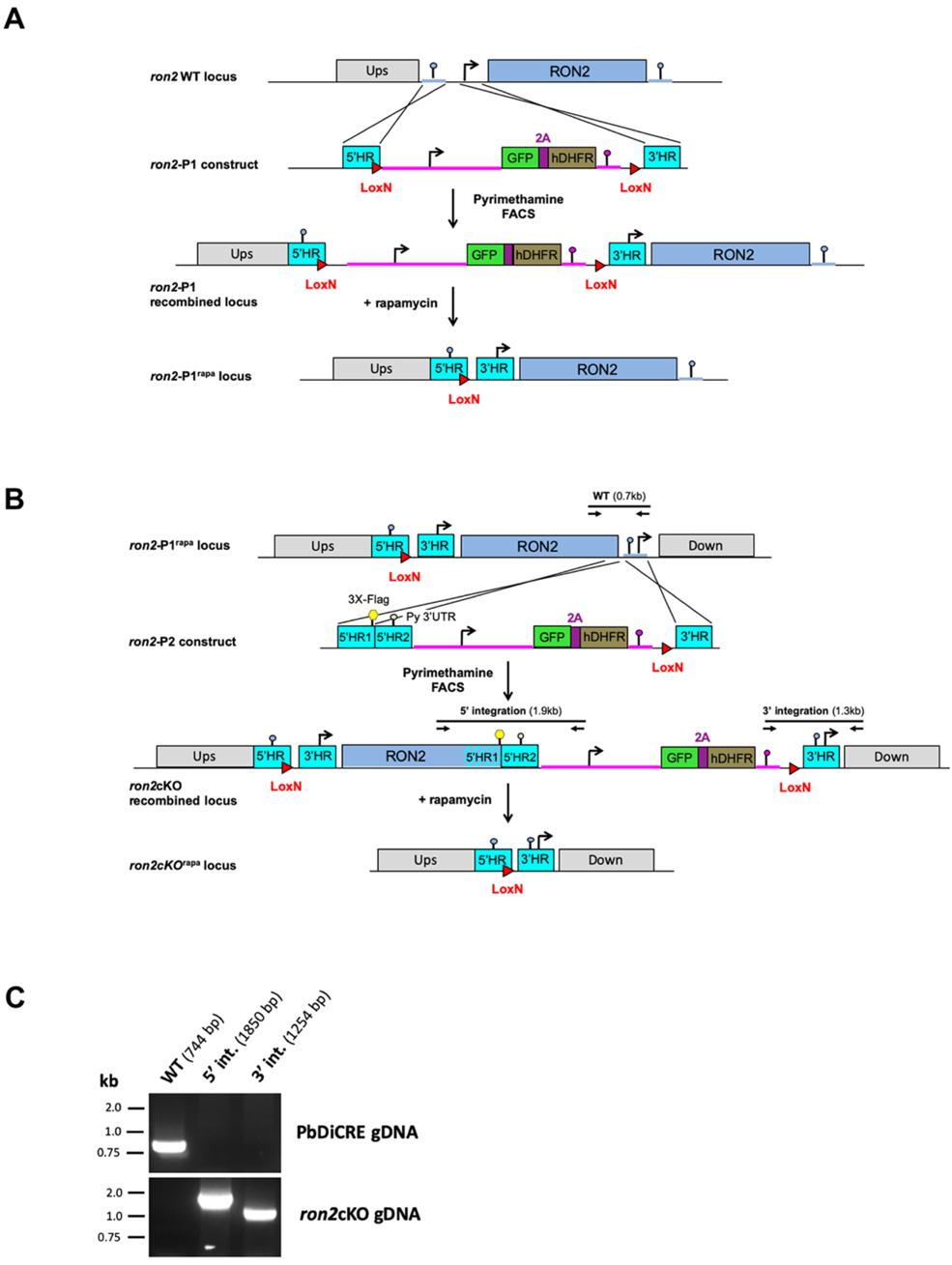
Generation of *ron2*cKO parasites using the DiCre system. **A-B.** Two-step strategy to generate *ron2*cKO parasites. In the first step (A), the *RON2* locus in PbDiCre parasites was targeted with a *ron2-*P1 replacement plasmid containing 5’ and 3’ homologous sequences and two LoxN sites flanking a GFP-2A-hDHFR cassette. Upon double crossover recombination, the two LoxN sites are inserted upstream of *RON2*. Activation of the DiCre recombinase with rapamycin results in excision of the GFP-2A-hDHFR cassette, leaving a single LoxN site upstream of the gene in *ron2*-P1^rapa^ parasites. In the second step (B), the *RON2* locus in *ron2*-P1^rapa^ parasites was targeted with a *ron2-*P2 replacement plasmid containing 5’ and 3’ homologous sequences flanking a GFP-2A-hDHFR cassette and a single LoxN site. Upon double crossover recombination, the LoxN site is inserted downstream of *RON2* and the GFP-2A-hDHFR cassette. Activation of the DiCre recombinase with rapamycin results in excision of the entire *RON2* gene together with the GFP-2A-hDHFR cassette. Genotyping primers and expected PCR fragments are indicated by arrows and lines, respectively. **C**. Genotyping of PbDiCre and *ron2*cKO parasites. Parasite genomic DNA was analysed by PCR using primer combinations specific for the unmodified locus (WT), the 5’ and 3’ integration events.

**Figure 4 – Supplement 2.**
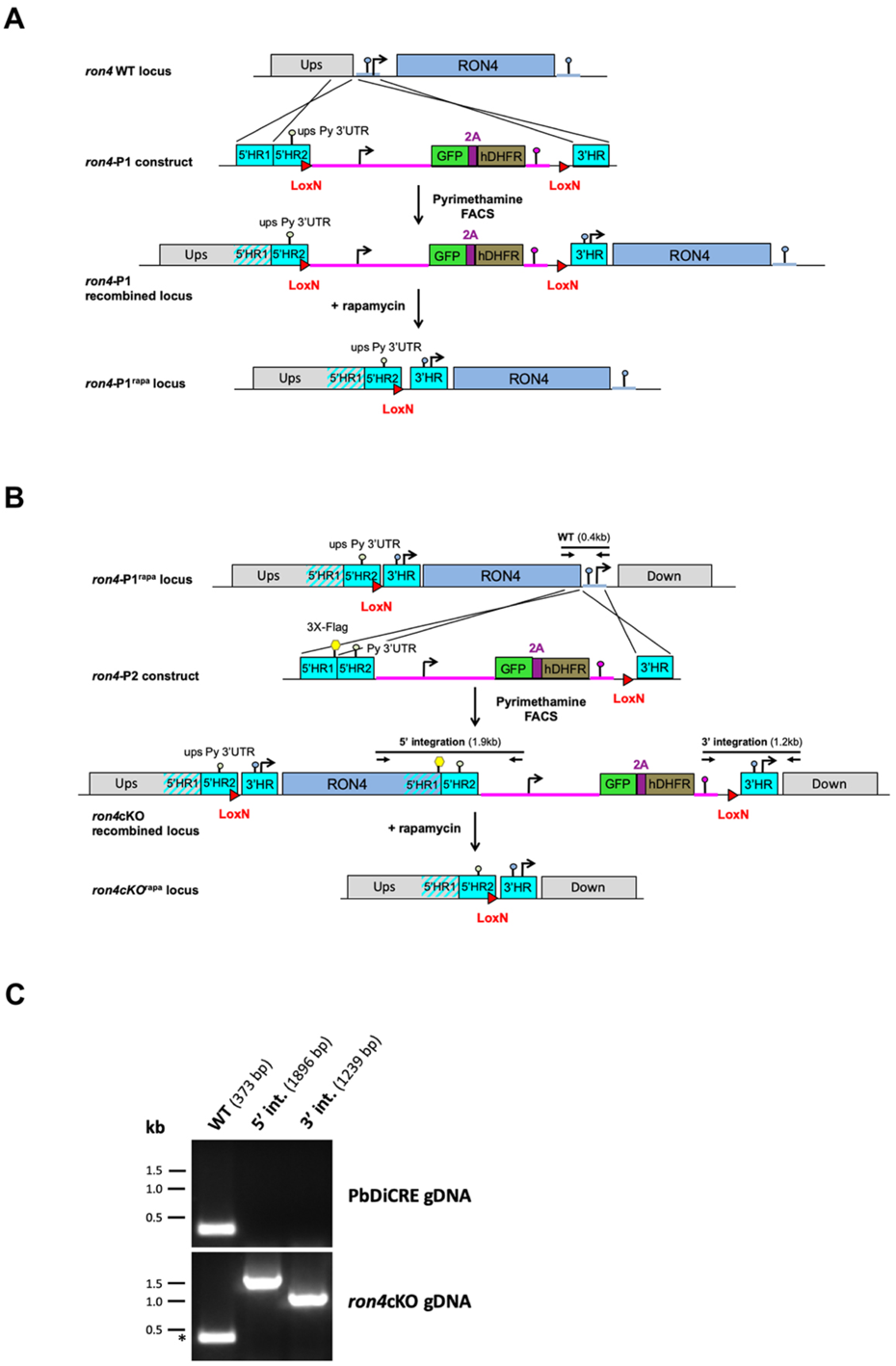
Generation of *ron2*cKO parasites using the DiCre system. **A-B.** Two-step strategy to generate *ron4*cKO parasites. In the first step (A), the *RON4* locus in PbDiCre parasites was targeted with a *ron2-*P1 replacement plasmid containing 5’ and 3’ homologous sequences and two LoxN sites flanking a GFP-2A-hDHFR cassette. Upon double crossover recombination, the two LoxN sites are inserted upstream of *RON4*. Activation of the DiCre recombinase with rapamycin results in excision of the GFP-2A-hDHFR cassette, leaving a single LoxN site upstream of the gene in *ron4*-P1^rapa^ parasites. In the second step (B), the *RON4* locus in *ron4*-P1^rapa^ parasites was targeted with a *ron4-*P2 replacement plasmid containing 5’ and 3’ homologous sequences flanking a GFP-2A-hDHFR cassette and a single LoxN site. Upon double crossover recombination, the LoxN site is inserted downstream of *RON4* and the GFP-2A-hDHFR cassette. Activation of the DiCre recombinase with rapamycin results in excision of the entire *RON4* gene together with the GFP-2A-hDHFR cassette. Genotyping primers and expected PCR fragments are indicated by arrows and lines, respectively. **C**. Genotyping of PbDiCre and *ron4*cKO parasites. Parasite genomic DNA was analysed by PCR using primer combinations specific for the unmodified locus (WT), the 5’ and 3’ integration events.

**Figure 4 – Supplement 3.**
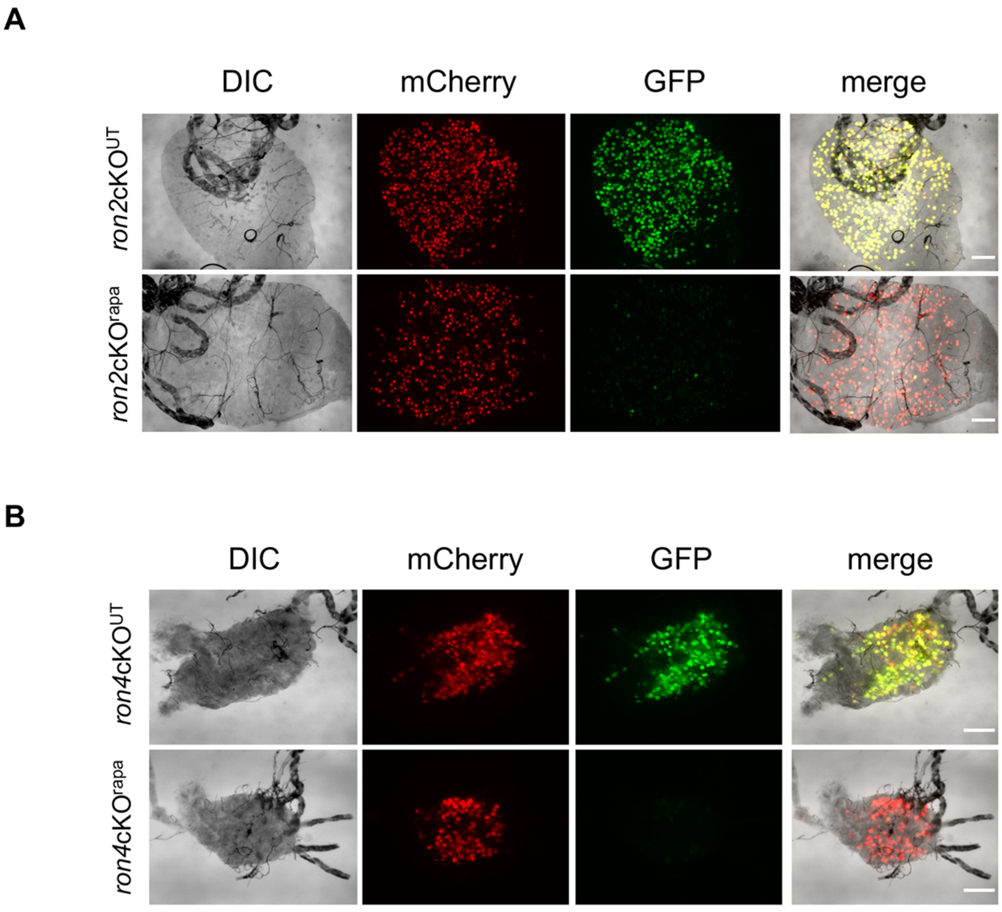
Imaging of *ron2*cKO and *ron4*cKO mosquito stages. **A-B.** Fluorescence microscopy of midguts from mosquitoes infected with untreated (UT) or rapamycin-treated (rapa) *ron2*cKO (A) or *ron4*cKO (B) parasites. Scale bar = 200 µm.

**Figure 4 – Supplement 4.**
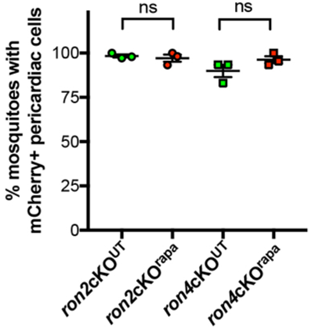
Analysis of mosquito pericardial structures. Quantification of mosquitoes with mCherry-labelled pericardial cells at D21 post-infection with untreated (UT) or rapamycin-treated (rapa) *ron2*cKO or *ron4*cKO parasites. Ns, non-significant (Two-tailed ratio paired t test).

**Figure 6 – Supplement 1.**
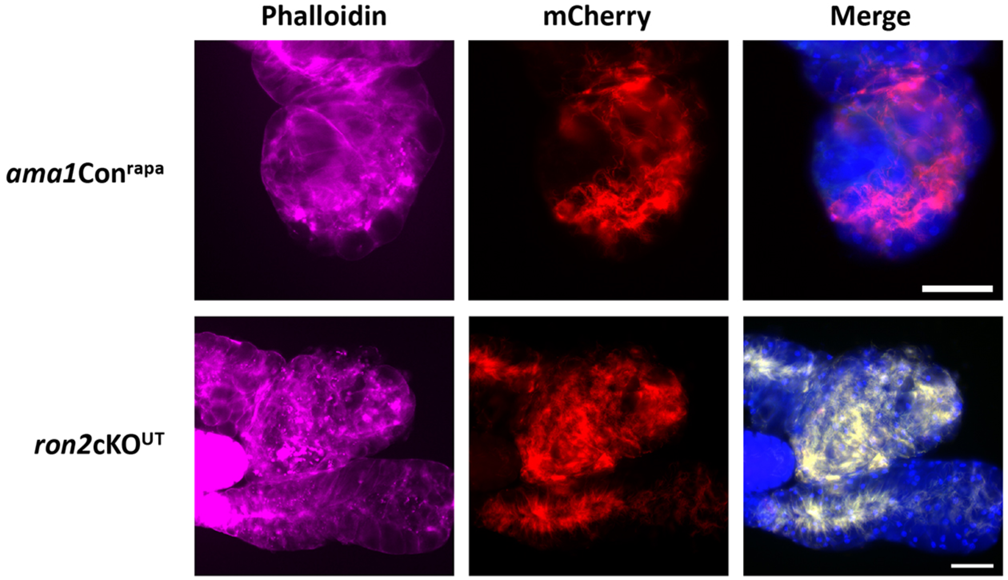
Fluorescence microscopy images of salivary gland distal lobes infected with rapamycin-treated *ama1*Con or untreated *ron2*cKO parasites. Samples were stained with Phalloidin-iFluor 647 (magenta) and Hoechst 77742 (Blue). The right panels show mCherry (red), GFP (green) and Hoechst (blue) merge images. In both cases, the heavy parasite load is associated with internal alterations of the phalloidin staining, but the basal border of the lobes is preserved. Scale bars, 50 μm.

**Supplementary Movie 1**. 3D segmentation of a mosquito salivary gland infected with WT sporozoites. Parasites appear in blue and secretory cavities in yellow. This movie corresponds to Figure 5A.

**Supplementary Movie 2**. 3D segmentation of a mosquito salivary gland infected with *ama1*cKO^rapa^ sporozoites. Parasites appear in blue and secretory cavities in yellow. This movie corresponds to Figure 5B.

**Supplementary Movie 3**. SBF-SEM sections of a mosquito salivary gland infected with WT parasites. This movie corresponds to Figure 5C-D.

## Supplementary tables

**Table S1**. Mass spectrometry analysis of co-IP from RON4-mCherry sporozoites.

**Table S2**. List of oligonucleotides used in the study.

**Table S3**. Quantitative data and statistical analysis.

## References

Bargieri DY, Andenmatten N, Lagal V, Thiberge S, Whitelaw JA, Tardieux I, Meissner M, Ménard R. 2013. Apical membrane antigen 1 mediates apicomplexan parasite attachment but is dispensable for host cell invasion. Nat Commun 4:2552. doi:10.1038/ncomms3552

Besteiro S, Dubremetz JF, Lebrun M. 2011. The moving junction of apicomplexan parasites: A key structure for invasion. Cell Microbiol 13:797–805. doi:10.1111/j.1462-5822.2011.01597.x

Bindschedler A, Wacker R, Egli J, Eickel N, Schmuckli-Maurer J, Franke-Fayard BM, Janse CJ, Heussler VT. 2020. Plasmodium berghei sporozoites in nonreplicative vacuole are eliminated by a PI3P-mediated autophagy-independent pathway. Cell Microbiol e13271. doi:10.1111/cmi.13271

Collins CR, Das S, Wong EH, Andenmatten N, Stallmach R, Hackett F, Herman JP, Müller S, Meissner M, Blackman MJ. 2013. Robust inducible Cre recombinase activity in the human malaria parasite Plasmodium falciparum enables efficient gene deletion within a single asexual erythrocytic growth cycle. Mol Microbiol 88:687–701. doi:10.1111/mmi.12206

Collins CR, Hackett F, Howell SA, Snijders AP, Russell MR, Collinson LM, Blackman MJ. 2020. The malaria parasite sheddase sub2 governs host red blood cell membrane sealing at invasion. Elife 9. doi:10.7554/ELIFE.61121

Cowman AF, Tonkin CJ, Tham W-H, Duraisingh MT. 2017. The Molecular Basis of Erythrocyte Invasion by Malaria Parasites. Cell Host Microbe 22:232–245. doi:10.1016/j.chom.2017.07.003

Deerinck TJ, Bushong E a., Thor a., Ellisman MH. 2010. NCMIR methods for 3D EM: A new protocol for preparation of biological specimens for serial block face scanning electron microscopy. Microscopy.

Ecker A, Lewis RE, Ekland EH, Jayabalasingham B, Fidock DA. 2012. Tricks in Plasmodium’s molecular repertoire - Escaping 3’UTR excision-based conditional silencing of the chloroquine resistance transporter gene. Int J Parasitol 42. doi:10.1016/j.ijpara.2012.09.003

Fernandes P, Briquet S, Patarot D, Loubens M, Hoareau-Coudert B, Silvie O. 2020. The dimerisable Cre recombinase allows conditional genome editing in the mosquito stages of Plasmodium berghei. PLoS One 15. doi:10.1371/journal.pone.0236616

Formaglio P, Tavares J, Ménard R, Amino R. 2014. Loss of host cell plasma membrane integrity following cell traversal by Plasmodium sporozoites in the skin. Parasitol Int 63:237–244. doi:10.1016/j.parint.2013.07.009

Frénal K, Dubremetz J-F, Lebrun M, Soldati-Favre D. 2017. Gliding motility powers invasion and egress in Apicomplexa. Nat Rev Microbiol 15:645–660. doi:10.1038/nrmicro.2017.86

Ghosh AK, Devenport M, Jethwaney D, Kalume DE, Pandey A, Anderson VE, Sultan AA, Kumar N, Jacobs-Lorena M. 2009. Malaria parasite invasion of the mosquito salivary gland requires interaction between the Plasmodium TRAP and the Anopheles saglin proteins. PLoS Pathog 5. doi:10.1371/journal.ppat.1000265

Giovannini D, Späth S, Lacroix C, Perazzi A, Bargieri D, Lagal V, Lebugle C, Combe A, Thiberge S, Baldacci P, Tardieux I, Ménard R. 2011. Independent roles of apical membrane antigen 1 and rhoptry neck proteins during host cell invasion by apicomplexa. Cell Host Microbe 10:591–602. doi:10.1016/j.chom.2011.10.012

Hamada S, Pionneau C, Parizot C, Silvie O, Chardonnet S, Marinach C. 2021. In-depth proteomic analysis of Plasmodium berghei sporozoites using trapped ion mobility spectrometry with parallel accumulation-serial fragmentation. Proteomics 21. doi:10.1002/pmic.202000305

Harris KS, Casey JL, Coley AM, Masciantonio R, Sabo JK, Keizer DW, Lee EF, McMahon A, Norton RS, Anders RF, Foley M. 2005. Binding hot spot for invasion inhibitory molecules on Plasmodium falciparum apical membrane antigen 1. Infect Immun 73. doi:10.1128/IAI.73.10.6981-6989.2005

Ishino T, Murata E, Tokunaga N, Baba M, Tachibana M, Thongkukiatkul A, Tsuboi T, Torii M. 2019. Rhoptry neck protein 2 expressed in Plasmodium sporozoites plays a crucial role during invasion of mosquito salivary glands. Cell Microbiol 21:e12964. doi:10.1111/cmi.12964

Janse CJ, Ramesar J, Waters AP. 2006. High-efficiency transfection and drug selection of genetically transformed blood stages of the rodent malaria parasite Plasmodium berghei. Nat Protoc 1:346–356. doi:10.1038/nprot.2006.53

Kappe SHI, Noe AR, Fraser TS, Blair PL, Adams JH. 1998. A family of chimeric erythrocyte binding proteins of malaria parasites. Proc Natl Acad Sci U S A 95. doi:10.1073/pnas.95.3.1230

Kariu T, Yuda M, Yano K, Chinzei Y. 2002. MAEBL is essential for malarial sporozoite infection of the mosquito salivary gland. J Exp Med 195:1317–1323. doi:10.1084/jem.20011876

Kudryashev M, Lepper S, Stanway R, Bohn S, Baumeister W, Cyrklaff M, Frischknecht F. 2010. Positioning of large organelles by a membrane-associated cytoskeleton in Plasmodium sporozoites. Cell Microbiol 12:362–371. doi:10.1111/j.1462-5822.2009.01399.x

Lamarque M, Besteiro S, Papoin J, Roques M, Vulliez-Le Normand B, Morlon-Guyot J, Dubremetz JF, Fauquenoy S, Tomavo S, Faber BW, Kocken CH, Thomas AW, Boulanger MJ, Bentley GA, Lebrun M. 2011. The RON2-AMA1 interaction is a critical step in moving junction-dependent invasion by apicomplexan parasites. PLoS Pathog 7. doi:10.1371/journal.ppat.1001276

Lamarque MH, Roques M, Kong-Hap M, Tonkin ML, Rugarabamu G, Marq J-B, Penarete-Vargas DM, Boulanger MJ, Soldati-Favre D, Lebrun M. 2014. Plasticity and redundancy among AMA-RON pairs ensure host cell entry of Toxoplasma parasites. Nat Commun 5:4098. doi:10.1038/ncomms5098

Lindner SE, Swearingen KE, Harupa A, Vaughan AM, Sinnis P, Moritz RL, Kappe SH. 2013. Total and putative surface proteomics of malaria parasite salivary gland sporozoites. Mol Cell Proteomics 12:1127–1143. doi:10.1074/mcp.M112.024505

Manzoni G, Briquet S, Risco-Castillo V. 2014. A rapid and robust selection procedure for generating drug-selectable marker-free recombinant malaria parasites. Sci Rep 99210:1–10. doi:10.1038/srep04760

Meis JFGM, Wismans PGP, Jap PHK, Lensen AHW, Ponnudurai T. 1992. A scanning electron microscopic study of the sporogonic development of Plasmodium falciparum in Anopheles stephensi. Acta Trop 50:227–236. doi:10.1016/0001-706X(92)90079-D

Mota MM, Pradel G, Vanderberg JP, Hafalla JC, Frevert U, Nussenzweig RS, Nussenzweig V, Rodriguez A. 2001. Migration of Plasmodium sporozoites through cells before infection. Science 291:141–144.

Nozaki M, Baba M, Tachibana M, Tokunaga N, Torii M, Ishino T. 2020. Detection of the Rhoptry Neck Protein Complex in Plasmodium Sporozoites and Its Contribution to Sporozoite Invasion of Salivary Glands. mSphere 5:e00325–20. doi:10.1128/msphere.00325-20

Pimenta PF, Touray M, Miller L. 1994. The Journey of Malaria Sporozoites in the Mosquito Salivary Gland. J Eukaryot Microbiol 41:608–624. doi:10.1111/j.1550-7408.1994.tb01523.x

Posthuma G, Meis JFGM, Verhave JP, Gigengack S, Hollingdale MR, Ponnudurai T, Geuze HJ. 1989. Immunogold determination of Plasmodium falciparum circumsporozoite protein in Anopheles stephensi salivary gland cells. Eur J Cell Biol 49.

Ramakrishnan C, Delves MJ, Lal K, Blagborough AM, Butcher G, Baker KW, Sinden RE. 2013. Laboratory maintenance of rodent malaria parasites. Methods Mol Biol 923:51– 72. doi:10.1007/978-1-62703-026-7_5

Richard D, MacRaild CA, Riglar DT, Chan JA, Foley M, Baum J, Ralph SA, Norton RS, Cowman AF. 2010. Interaction between Plasmodium falciparum apical membrane antigen 1 and the rhoptry neck protein complex defines a key step in the erythrocyte invasion process of malaria parasites. J Biol Chem 285. doi:10.1074/jbc.M109.080770

Riglar DT, Richard D, Wilson DW, Boyle MJ, Dekiwadia C, Turnbull L, Angrisano F, Marapana DS, Rogers KL, Whitchurch CB, Beeson JG, Cowman AF, Ralph SA, Baum J. 2011. Super-resolution dissection of coordinated events during malaria parasite invasion of the human erythrocyte. Cell Host Microbe 9:9–20. doi:10.1016/j.chom.2010.12.003

Risco-Castillo V. 2015. Malaria sporozoites traverse cells within transient vacuoles. Cell Host Microbe.

Risco-Castillo V, Topçu S, Marinach C, Manzoni G, Bigorgne AE, Briquet S, Baudin X, Lebrun M, Dubremetz JF, Silvie O. 2015. Malaria sporozoites traverse host cells within transient vacuoles. Cell Host Microbe 18:593–603. doi:10.1016/j.chom.2015.10.006

Saenz FE, Balu B, Smith J, Mendonca SR, Adams JH. 2008. The transmembrane isoform of Plasmodium falciparum MAEBL is essential for the invasion of Anopheles salivary glands. PLoS One 3:e2287. doi:10.1371/journal.pone.0002287

Schrevel J, Asfaux-Foucher G, Hopkins JM, Robert V, Bourgouin C, Prensier G, Bannister LH. 2008. Vesicle trafficking during sporozoite development in Plasmodium berghei: Ultrastructural evidence for a novel trafficking mechanism. Parasitology 135. doi:10.1017/S0031182007003629

Silvie O, Franetich J-FFJ-F, Charrin S, Mueller MSSMS, Siau A, Bodescot M, Rubinstein E, Hannoun L, Charoenvit Y, Kocken CHHCH, Thomas AWWAW, Van Gemert G-JG-JJ, Sauerwein RWRWW, Blackman MJJMJ, Anders RFRFF, Pluschke G, Mazier D. 2004. A role for apical membrane antigen 1 during invasion of hepatocytes by Plasmodium falciparum sporozoites. J Biol Chem 279:9490–9496. doi:10.1074/jbc.M311331200

Silvie O, Franetich JF, Boucheix C, Rubinstein E, Mazier D. 2007. Alternative invasion pathways for plasmodium berghei sporozoites. Int J Parasitol 37:173–182. doi:10.1016/j.ijpara.2006.10.005

Sinden RE, Strong K. 1978. An ultrastructural study of the sporogonic development of plasmodium falciparum in anopheles gambiae. Trans R Soc Trop Med Hyg 72:477–491. doi:10.1016/0035-9203(78)90167-0

Srinivasan P, Baldeviano GC, Miura K, Diouf A, Ventocilla JA, Leiva KP, Lugo-Roman L, Lucas C, Orr-Gonzalez S, Zhu D, Villasante E, Soisson L, Narum DL, Pierce SK, Long CA, Diggs C, Duffy PE, Lescano AG, Miller LH. 2017. A malaria vaccine protects Aotus monkeys against virulent Plasmodium falciparum infection. npj Vaccines 2. doi:10.1038/s41541-017-0015-7

Srinivasan P, Beatty WL, Diouf A, Herrera R, Ambroggio X, Moch JK, Tyler JS, Narum DL, Pierce SK, Boothroyd JC, Haynese JD, Millera LH. 2011. Binding of Plasmodium merozoite proteins RON2 and AMA1 triggers commitment to invasion. Proc Natl Acad Sci U S A 108. doi:10.1073/pnas.1110303108

Srinivasan P, Ekanem E, Diouf A, Tonkin ML, Miura K, Boulanger MJ, Long CA, Narum DL, Miller LH. 2014. Immunization with a functional protein complex required for erythrocyte invasion protects against lethal malaria. Proc Natl Acad Sci U S A 111. doi:10.1073/pnas.1409928111

Sterling CR, Aikawa M, Vanderberg JP. 1973. The passage of Plasmodium berghei sporozoites through the salivary glands of Anopheles stephensi: An electron microscope study. J Parasitol 59. doi:10.2307/3278847

Swearingen KE, Lindner SE, Flannery EL, Vaughan AM, Morrison RD, Patrapuvich R, Koepfli C, Muller I, Jex A, Moritz RL, Kappe SHI, Sattabongkot J, Mikolajczak SA. 2017. Proteogenomic analysis of the total and surface-exposed proteomes of Plasmodium vivax salivary gland sporozoites. PLoS Negl Trop Dis 11:e0005791. doi:10.1371/journal.pntd.0005791

Tibúrcio M, Yang ASP, Yahata K, Suárez-Cortés P, Belda H, Baumgarten S, van de Vegte-Bolmer M, van Gemert G-J, van Waardenburg Y, Levashina EA, Sauerwein RW, Treeck M. 2019. A Novel Tool for the Generation of Conditional Knockouts To Study Gene Function across the Plasmodium falciparum Life Cycle. MBio 10:e01170–19. doi:10.1128/mbio.01170-19

Tokunaga N, Nozaki M, Tachibana M, Baba M, Matsuoka K, Tsuboi T, Torii M, Ishino T. 2019. Expression and Localization Profiles of Rhoptry Proteins in Plasmodium berghei Sporozoites. Front Cell Infect Microbiol 9:316. doi:10.3389/fcimb.2019.00316

Tufet-Bayona M, Janse CJ, Khan SM, Waters AP, Sinden RE, Franke-Fayard B. 2009. Localisation and timing of expression of putative Plasmodium berghei rhoptry proteins in merozoites and sporozoites. Mol Biochem Parasitol 166:22–31.

Walton J. 1979. Lead aspartate, an en bloc contrast stain particularly useful for ultrastructural enzymology. J Histochem Cytochem 27. doi:10.1177/27.10.512319

Wells MB, Andrew DJ. 2019. Anopheles salivary gland architecture shapes plasmodium sporozoite availability for transmission. MBio 10. doi:10.1128/mBio.01238-19

Yang ASP, Lopaticki S, O’Neill MT, Erickson SM, Douglas DN, Kneteman NM, Boddey JA. 2017. AMA1 and MAEBL are important for Plasmodium falciparum sporozoite infection of the liver. Cell Microbiol 19:e12745. doi:10.1111/cmi.12745

